# Multiple pathways for reestablishing PAR polarity in *C. elegans* embryo

**DOI:** 10.1101/2022.12.15.520651

**Authors:** Laurel A. Koch, Lesilee S. Rose

**Affiliations:** Department of Molecular and Cellular Biology and Integrative Genetics and Genomics Graduate Program, University of California, Davis

**Keywords:** asymmetric division, polarization, asymmetry, embryo

## Abstract

Asymmetric cell divisions, where cells divide with respect to a polarized axis and give rise to daughter cells with different fates, are critically important during development. In many such divisions, the conserved PAR polarity proteins accumulate in distinct cortical domains in response to a symmetry breaking cue. The one-cell *C. elegans* embryo has been a paradigm for understanding mechanisms of PAR polarization, but much less is known about polarity reestablishment during subsequent divisions. Here, we investigate the polarization of the P1 cell of the two-cell embryo. A posterior PAR-2 domain forms in the first four minutes of the birth of P1, and polarization becomes stronger over time. Initial polarization depends on the PKC-3 and PAR-1 kinases. However, in *par-1* mutants, delayed polarization can occur, at a time when centrosome-associated AIR-1 is near the posterior cortex and myosin flows towards the anterior. Loss of myosin and *par-1* function together results in more severe polarity defects. Based on these and other results, we propose that PAR polarity in the P1 cell is generated by at least two redundant mechanisms: There is a novel early pathway dependent on PAR-1, PKC-3 and cytoplasmic polarity, and a late pathway that resembles symmetry breaking in the one-cell embryo and requires myosin flow and PKC-3.

## Introduction

Asymmetric cell division is the process in which one cell divides to give rise to two daughter cells with different cell fates. This process is important for generating cell diversity throughout development, as well as for maintaining stem cell populations in many organisms. One of the important steps in many asymmetric cell divisions is generating a polarity axis that will be bisected by cytokinesis, so that cell fate determinants are segregated differentially to the daughter cells (Sunchu & Cabernard, 2020; Venkei & Yamashita, 2018). In many organisms, the cortically localized partitioning-defective (PAR) proteins are required for generating this polarity axis. For example, the PAR proteins regulate asymmetric cell division in the *C. elegans* one-cell embryo and the Drosophila neuroblast. In these and other systems, groups of PAR proteins become localized to reciprocal, mutually exclusive cortical domains, for example anteriorly localized PARs (aPARs) and posteriorly localized PARs (pPARs)(Goldstein & Macara, 2007; Pickett et al., 2019; Rose & Gonczy, 2014).

The initial establishment of PAR polarity domains occurs in response to various symmetry breaking cues. In the one-cell *C. elegans* embryo (called P0), symmetry is broken when the sperm fertilizes the oocyte (Cowan & Hyman, 2004). At the time of fertilization, the aPAR proteins, PAR-3 and PAR-6, which are PDZ domain containing scaffolding proteins, and PKC-3, which is an atypical kinase C, are uniform on the cortex (Cuenca et al., 2003; Kemphues et al., 1988; Tabuse et al., 1998; Watts et al., 1996). The sperm derived centrioles recruit centrosome components including Aurora A Kinase (AIR-1), which locally inhibits actomyosin contractility, generating actomyosin flow away from the location of the centrosome. This flow moves the aPARs towards the opposite end of the embryo, which will become the anterior pole (Klinkert et al., 2019; Munro et al., 2004; Schonegg et al., 2014; Zhao et al., 2019). As the aPARs clear from the posterior this allows PAR-1, a serine/threonine kinase, and PAR-2, a RING domain protein, to move onto the posterior cortex forming the pPAR domain. (Boyd et al., 1996; Cheeks et al., 2004; Cowan & Hyman, 2004; Cuenca et al., 2003; Guo & Kemphues, 1995; Hao et al., 2006). In this and other systems, the reciprocal PAR domains are then maintained by mutual exclusion. PKC-3 phosphorylates PAR-1 and PAR-2, restricting the pPARs from the anterior. Meanwhile, PAR-1 phosphorylates PAR-3 to inhibit PAR-3 association with the cortex, which with other mechanisms in *C. elegans* restricts aPARs from the posterior (Benton & St Johnston, 2003; Hao et al., 2006; Hurov et al., 2004).

There is also a redundant pathway that can establish PAR polarity in the P0 cell when actomyosin flow is inhibited. In this backup pathway, PAR-2 appears to establish polarity by binding to microtubules emanating from the sperm-derived centrosomes. Microtubule association shelters PAR-2 from being phosphorylated by PKC-3, allowing PAR-2 to accumulate on the posterior cortex and then PAR-1 can load onto the cortex (Motegi et al., 2011; Zonies et al., 2010). This pathway thus generates the same reciprocal aPAR and pPAR domains, but with a temporal delay.

PAR domains are important for correctly segregating cytoplasmic polarity and orienting the spindle, so that the P0 cell divides to give rise to a larger AB cell and smaller P1 cell with different fates and division patterns. The PAR polarity axis is reestablished in P1 and every subsequent germ-line P cell, which all divide asymmetrically (Rose & Gonczy, 2014). After P0 cytokinesis, the P1 cell has PAR-2 around the entire cortex. In response to an unknown polarity cue, an anterior-posterior PAR polarity axis forms again. At the four-cell stage, the P2 cell also inherits PAR-2 uniformly; aPAR and pPAR domains form in this cell, oriented by MES-1/SRC-1signalling at the EMS/P2 cell contact (Arata et al., 2010; Bei et al., 2002). In contrast, contact between the AB and P1 cell is not required for the spindle to rotate or divide asymmetrically (Goldstein, 1993, 1995). However, previous work has shown that there is actomyosin flow in the P1 cell towards the anterior and this correlates with the movement of PAR-6 on the cortex (Munro et al., 2004). The P1 cell nucleus also moves posteriorly after the end of P0 cytokinesis, making it plausible that the centrosome and AIR-1 might participate in the repolarization of P1, in a mechanism similar to that used in the P0 cell.

In this study, we sought to understand how PAR polarity is reestablished in the P1 cell. Here we show that the PAR-2 domain starts to form in the P1 cell within two minutes of the end of P0 cytokinesis, before nuclear movement towards the posterior or actomyosin flow occurs. We also found that *par-1* mutants exhibit late polarization, and defects in polarization correlate with incorrect cytoplasmic partitioning in the P0 cell. Our findings support a model whereby theP1 cell polarizes via at least two redundant pathways. The early pathway requires proper cytoplasmic polarity, PAR-1 and PKC-3, and clears PAR-2 from the anterior to form a PAR-2 domain. The late pathway utilizes actomyosin flow and causes aPAR accumulation in the anterior, creating a more strongly polarized distribution of PAR-2. These results will further our understanding of how polarity can be established in response to different cues.

## Results

### Determining the timing of PAR polarization in the P1 cell

To determine what components are present at the birth of the P1 cell and the timing of P1 cell polarization, we characterized the formation of PAR polarity using embryos co-labeled with endogenously tagged mCh::PAR-2 and PAR-6::GFP (Reich et al., 2019). Embryos were imaged from the end of P0 cytokinesis (time 0:00) through the end of the P1 cell cycle, and the cortical localization of PAR-2 and PAR-6 were quantified throughout the first ten minutes of the cell cycle (Fig 1A). During cytokinesis PAR-2 moved in with the furrow and only appeared to be present on the P1 side of the furrow (Fig. S1). Thus, at the birth of the P1 cell PAR-2 was localized around the entire cortex as previously described (Cuenca et al., 2003), and quantification showed that the levels are uniform on average (Fig. 1B, C). In some embryos, the levels of PAR-2 at the AB-P1 cell contact started to decrease as early as one minute after P0 cell cytokinesis, and in all embryos PAR-2 levels decreased at the AB-P1 cell contact by two minutes. We refer to this decrease as “clearing of PAR-2” and note that clearing was also accompanied by a corresponding increase of PAR-2 at the posterior cortex of the P1 cell. PAR-2 continues to clear from the anterior until 6 minutes and continues to accumulate in the posterior for the whole cell cycle (Fig. 1B). As a way to quantify the initial clearing of PAR-2 versus the formation of a more discrete posterior domain over time in this and mutant backgrounds, we measured the posterior/anterior ratio of PAR-2 at four and eight minutes after cytokinesis. At four minutes the posterior levels were three times higher than the anterior, and at eight minutes the posterior levels were seven times higher (Fig. 1D, E). These data indicate that the P1 cell is polarized with respect to PAR-2 by as early as four minutes, and the posterior domain of PAR-2 becomes stronger as the cell cycle continues.

**Figure 1.**
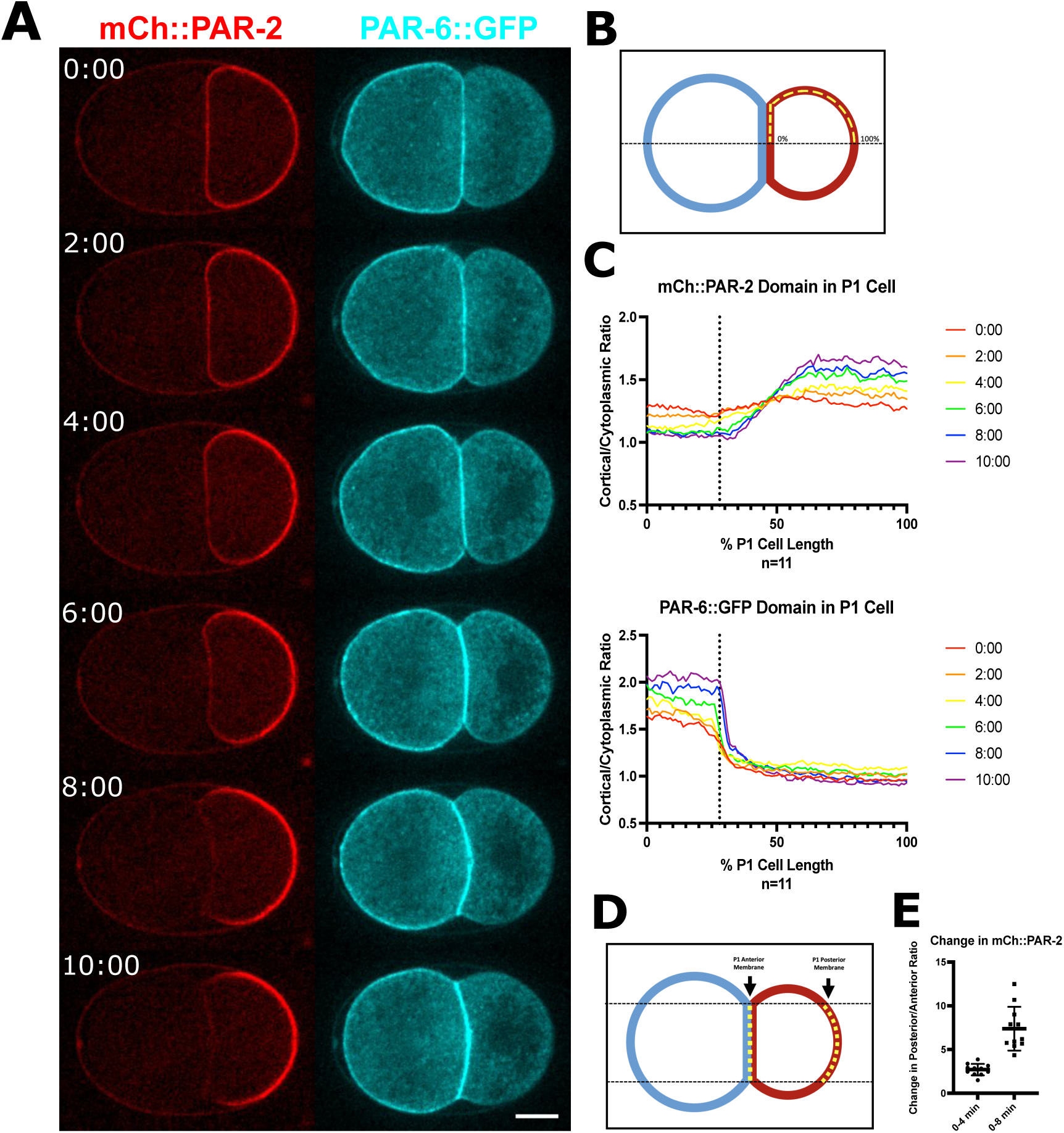
P1 cell polarization starts within four minutes of cytokinesis. (A) Fluorescent images of an embryo expressing mCh::PAR-2 and GFP::PAR-6. Anterior is to the left and posterior to the right in this and all figures. Time zero equals completion of cytokinesis. Scale bar is 10um. See also Supplemental Video 1. (B) Illustration of quantification of cortical fluorescence. Embryos were divided along the AP axis; the cortex was traced in Image J (yellow dotted line) from the cell contact (0%) to the posterior (100%) and then divided by cytoplasmic intensity. (C) Fluorescence intensity ratio plots every two minutes for the first ten minutes of the P1 cell cycle; each line is an average of multiple embryos. Dotted line highlights the end of the AB-P1 cell contact. (D) Illustration of how change in fluorescence measurements were taken for each time point. (E) Quantification of the change in posterior/anterior ratio from 0 to 4 minutes and 0 to 8 minutes after cytokinesis in control embryos. Means and statistics are reported in Supplemental Table 2.

Similar to PAR-2, PAR-6 signal moved in on the cytokinetic furrow but appeared to be present mainly on the AB side (Fig. S1). At cytokinesis completion, PAR-6 signal appeared highest at the AB-P1 cell contact and was at low levels throughout the P1 cell. For the first four minutes after cytokinesis, PAR-6 cortical levels increased globally. After four minutes, PAR-6 levels started to go down in the posterior and continued to accumulate in the anterior (Fig 1B). We cannot distinguish whether the increase in signal at the cell contact is in the AB or P1 cell. However, given that posterior levels of PAR-6 only start to decrease at four minutes, these data suggest that the polarization of PAR-6 may occur later than that of PAR-2.

### Early PAR-2 clearing does not correlate with AIR-1 localization and actomyosin flow

To test the hypothesis that AIR-1 on the centrosome is acting as the symmetry breaking cue in the P1 cell, we first looked at where the nucleus and centrosome were located during polarization. If centrosomal AIR-1 is a localized cue as in the one-cell embryo, we predicted that the nucleus and centrosome would move close to the posterior before the clearing of PAR-2 from the AB-P1 cell contact. We thus imaged embryos in DIC and measured the closest distance achieved between the posterior edge of the P1 nucleus and the posterior cortex (Fig 2A). On average, the nucleus was 24.5% of P1 cell length from the posterior, and the P1 nucleus reached its most posterior point at 5.15 minutes after P0 cytokinesis. The timing of movement was highly variable (SD=1.12 minutes) (Fig. 2B, C). Because PAR-2 polarization occurs before posterior nuclear movement, these data do not support the idea that proximity of the nuclear-centrosome complex to the cortex causes polarization.

**Figure 2.**
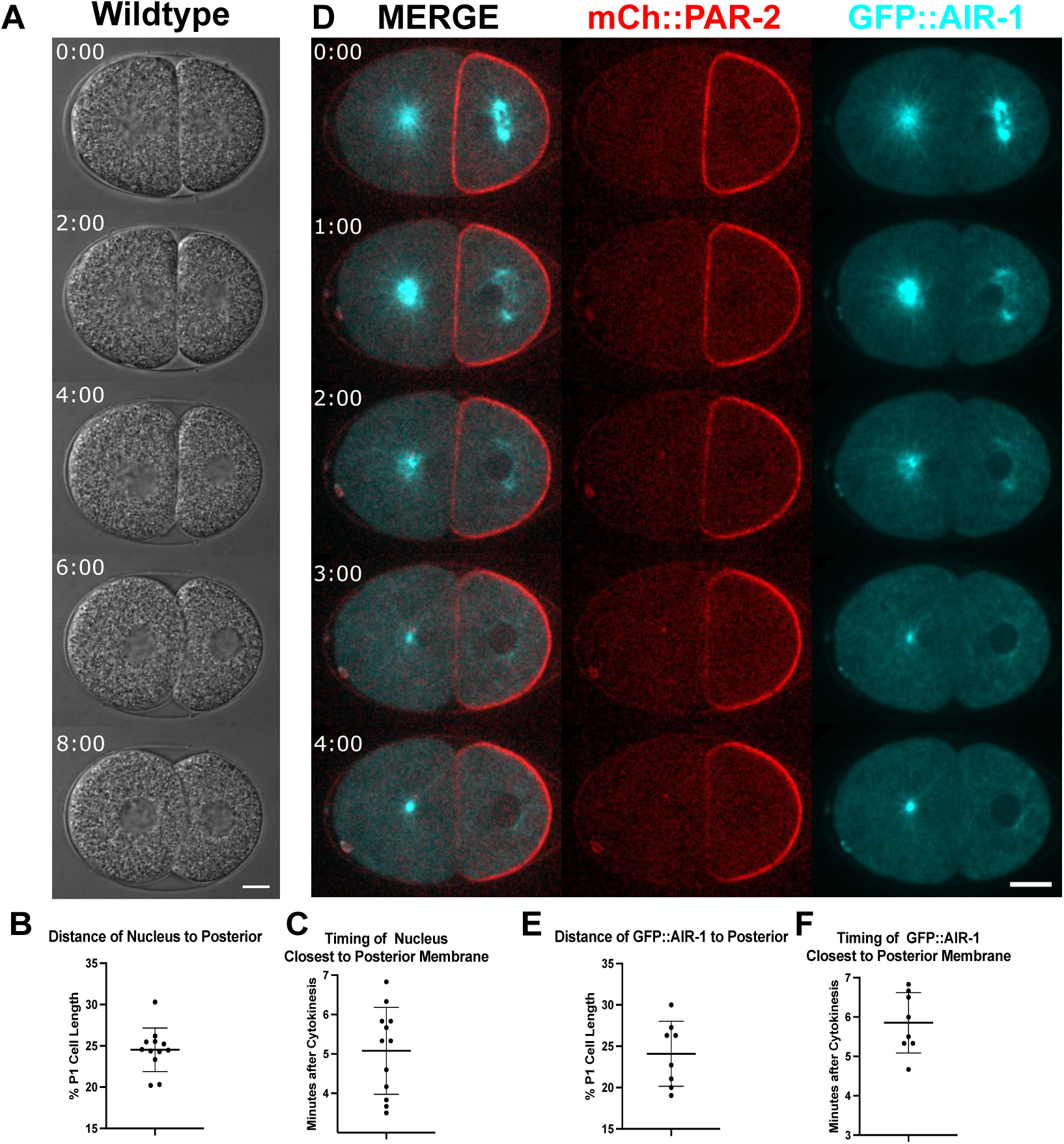
Nuclear-centrosome movement in the P1 cell does not correlate with early P1 polarization. (A) DIC images of nuclear movement in the P1 cell in wild-type embryo. Scale bar is 10um. See also Supplemental Video 2. (B) Quantification of the closest distance measured between the P1 nucleus and the posterior cortex. (C) Quantification of the time after cytokinesis when the nucleus is closest to the P1 cell posterior membrane (n=12). The total length of the P1 cell cycle is 16.09 minutes (N=11)(Fig. S2). (D) Confocal fluorescent images of embryos expressing mCh::PAR-2 and GFP::AIR-1 every minute from 0-4 minutes after the completion of cytokinesis. Scale bar is 10um. See also Supplemental Video 3. (E) Quantification of the closest distance of GFP::AIR-1 foci to the posterior membrane. (F) Quantification of the time after cytokinesis when GFP::AIR-1 foci is closest to the P1 cell posterior membrane (n=8). Means and statistics are reported in Supplemental Table 2.

To visualize the position of the centrosome specifically, we examined embryos expressing mCh::PAR-2 and GFP::AIR-1 (Fig 2C) (Portier et al., 2007). At the end of P0 cytokinesis, AIR-1 was localized on the posterior centrosome of the spindle, which at this stage has a disk shape in the center of the cell. AIR-1 was localized in a diffuse cloud around the disk aster during the first two minutes of the P1 cell cycle and then dissociated from the centrosome by three minutes. Around four minutes, AIR-1 started to accumulate onto the new maturing centrosomes. We found that on average, AIR-1 on the centrosome was closest to the posterior (24.09% P1 cell length) at 5.85 minutes after cytokinesis (Fig. 2D). These results do not support AIR-1 being the symmetry breaking cue in the P1 cell, because it is not posteriorly localized until 1.8 min after the time of PAR-2 clearing.

Previous studies also showed that there is anterior-directed actomyosin flow in the P1 cell which correlates with the movement of PAR-6 to the anterior. To better understand the timing and role of this flow we imaged embryos expressing GFP::NMY-2 on the cortex, and we also imaged GFP::NMY-2 in a mid-focal plane with mCh::PAR-2 (Fig. 3A, S3A-B) (Zonies et al., 2010). In some embryos (4/7), some anterior-directed flow of NMY-2 was visible for the first 1-2 minutes, but then ceased; this flow likely reflects the actomyosin movements of cytokinesis. A strong flow of GFP::NMY-2, similar to what was previously reported (Munro et al., 2004) started 4-6 minutes after cytokinesis completion; flow continued through at least ten minutes, and an anterior domain of GFP::NMY-2 became visible both cortically and in mid-plane (Fig 3A). These data show that NMY-2 flow, which is known to correlate with PAR-6 movement, does not occur until after the PAR-2 has already cleared substantially from the anterior of the P1 cell at 4 min.

**Figure 3.**
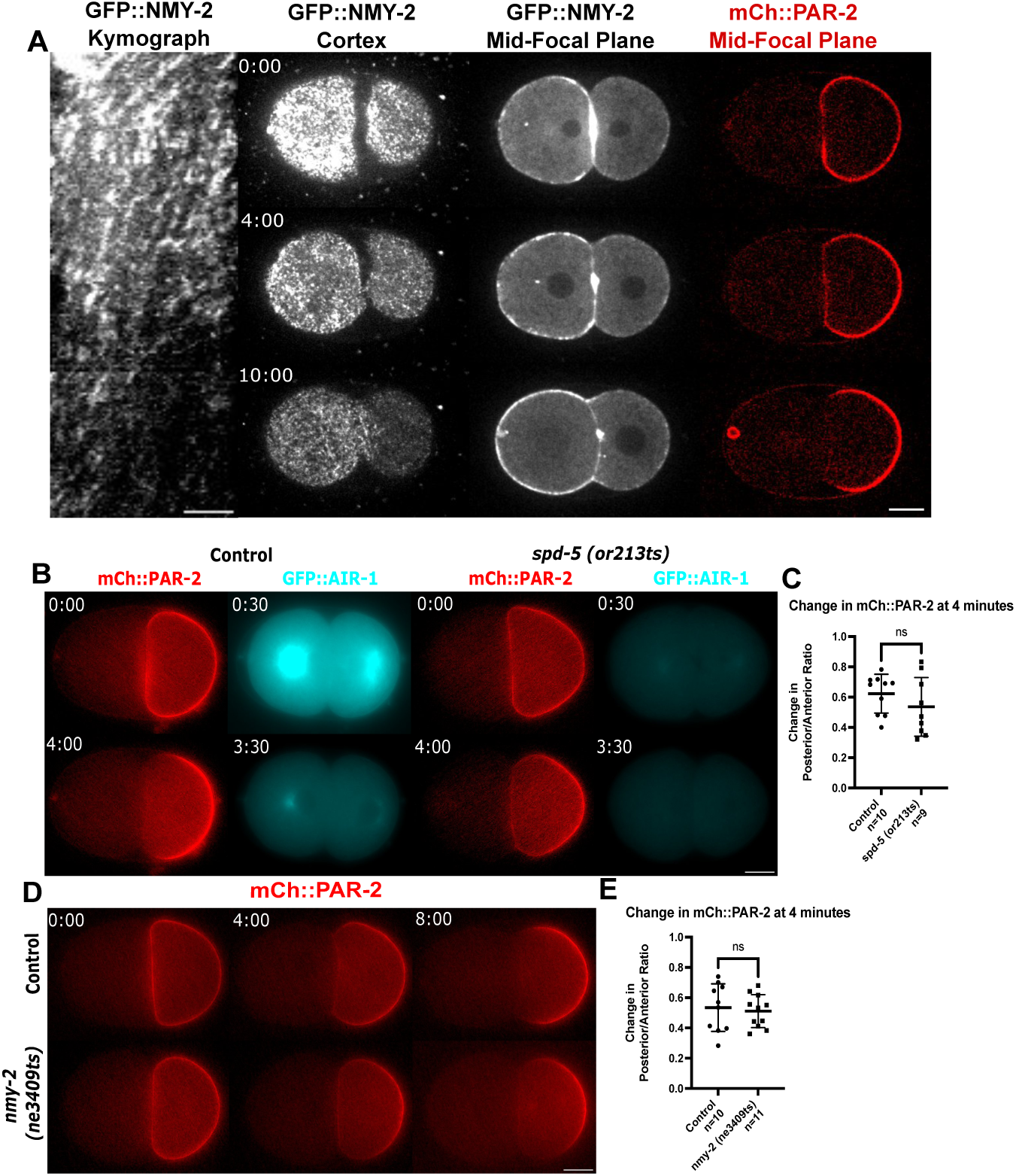
Actomyosin flow does not correlate with early P1 cell polarization. (A) Left: Kymograph of GFP::NMY-2 taken across the longest anterior-posterior axis of the P1 cell in the embryo shown in next panel . Scale bar is 5um. Second panel: Confocal fluorescent images of embryo expressing GFP::NMY-2 (Myosin) from a surface view. See also Supplemental Video 4. Third and fourth panels: Confocal fluorescent images of an embryo expressing GFP::NMY-2 and mCh::PAR-2 from a mid-focal plane. Scale bar is 10um. (B) Epiflourescent images of embryos expressing mCh::PAR-2 and GFP::AIR-1 with or without *spd-5 (or213ts);* embryos were shifted from 16C to 26C at P0 cell NEB. Scale bar is 10um. (C) Quantification of the change in posterior/anterior ratio from 0 to 4 minutes after cytokinesis in control and *spd-5 (or213ts)* embryos. (D) Epiflourescent images of embryos expressing mCh::PAR-2 with or without *nmy-2 (ne3409ts),* embryos were shifted from 16C to 26C at the end of P0 cytokinesis. Scale bar is 10um. (E) Quantification of the change in posterior/anterior ratio from 0 to 4 minutes after cytokinesis in control and *nmy-2 (ne3409ts)* embryos. Note that specific AP ratio values differ from those in other figures where confocal images were measured (see Methods). Means and statistics are reported in Supplemental Table 2.

Although AIR-1 does not seem to be localized near the posterior at the time of polarization, we were still interested in whether it could play a role in P1 cell polarization. AIR-1 is required in the P0 cell for normal polarization and cytokinesis, and conditional alleles are not available. Thus, to reduce AIR-1 levels on centrosomes before PAR-2 polarization, we used a fast-inactivating temperature sensitive mutant, *spd-5(or213ts).* SPD-5 is a centrosome maturation factor required to recruit AIR-1 to the centrosome (Hamill et al., 2002). Embryos were shifted to the restrictive temperature (26°C) during P0 cell NEB, and mCh::PAR-2 and GFP::AIR-1 were examined. By the end of P0 cytokinesis, AIR-1 levels were greatly reduced on the centrosome (Fig. 3B). The increase in the posterior/anterior ratio of mCh::PAR-2 between the end of cytokinesis and four minutes was similar for controls and *spd-5 (or213ts)* embryos (Fig. 3C). This result, along with AIR-1’s localization at the time of polarity establishment, leads us to believe that AIR-1 on centrosomes is not acting as the symmetry breaking cue in the P1 cell.

To directly test the role of actomyosin flow in the P1 cell, we used a fast-inactivating temperature sensitive mutant *nmy-2(ne3409ts)* and imaged mCh::PAR-2 (Fig. 3B) (Fievet et al., 2013; Liu et al., 2010). NMY-2 is required for cytokinesis, and shifting embryos at the start of furrow ingression led to a failure of cytokinesis; such shifts indicated that NMY-2 function was affected within two minutes of the shift to restrictive temperature. To make sure cytokinesis completed and the P1 cell inherited uniform PAR-2, *nmy-2(ne3409ts)* embryos were shifted to a restrictive temperature (26°C) at the end of cytokinesis. We found that there was no difference in the timing of PAR-2 clearing compared to controls shifted in the same way (Fig. 3D, E). These results are consistent with the view that NMY-2 flows do not stimulate early symmetry breaking in the P1 cell.

### PAR-1 and PKC-3 are inherited asymmetrically in the P1 cell

Since the mechanism of polarity establishment in the P1 cell appears to be different than that of the P0 cell, we examined whether other PAR proteins might be inherited asymmetrically and serve as a cue for P1 repolarization (Fig. 4A). PAR-1 is known to form both a cortical and cytoplasmic gradient in the P0 cell, but its localization in the P1 has not been examined in detail (Griffin et al., 2011). During furrowing, endogenously tagged PAR-1::meGFP appeared to only move into the furrow on the P1 side (Fig. S4A). At the end of cytokinesis, PAR-1::meGFP was present on part of the AB-P1 cell contact. However, PAR-1 was absent from the anterior corners of the P1 cell adjacent the contact, and levels increased towards the posterior. Similar results were seen with transgenic PAR-1::GFP (Fig. S4B-C). Thus, unlike PAR-2, cortical PAR-1 was inherited in a posterior cortical gradient (Fig. 4B). Similar to PAR-2, PAR-1 cleared from the contact within two minutes and became more enriched in the posterior cortex during the cell cycle (Fig. 4C).

**Figure 4.**
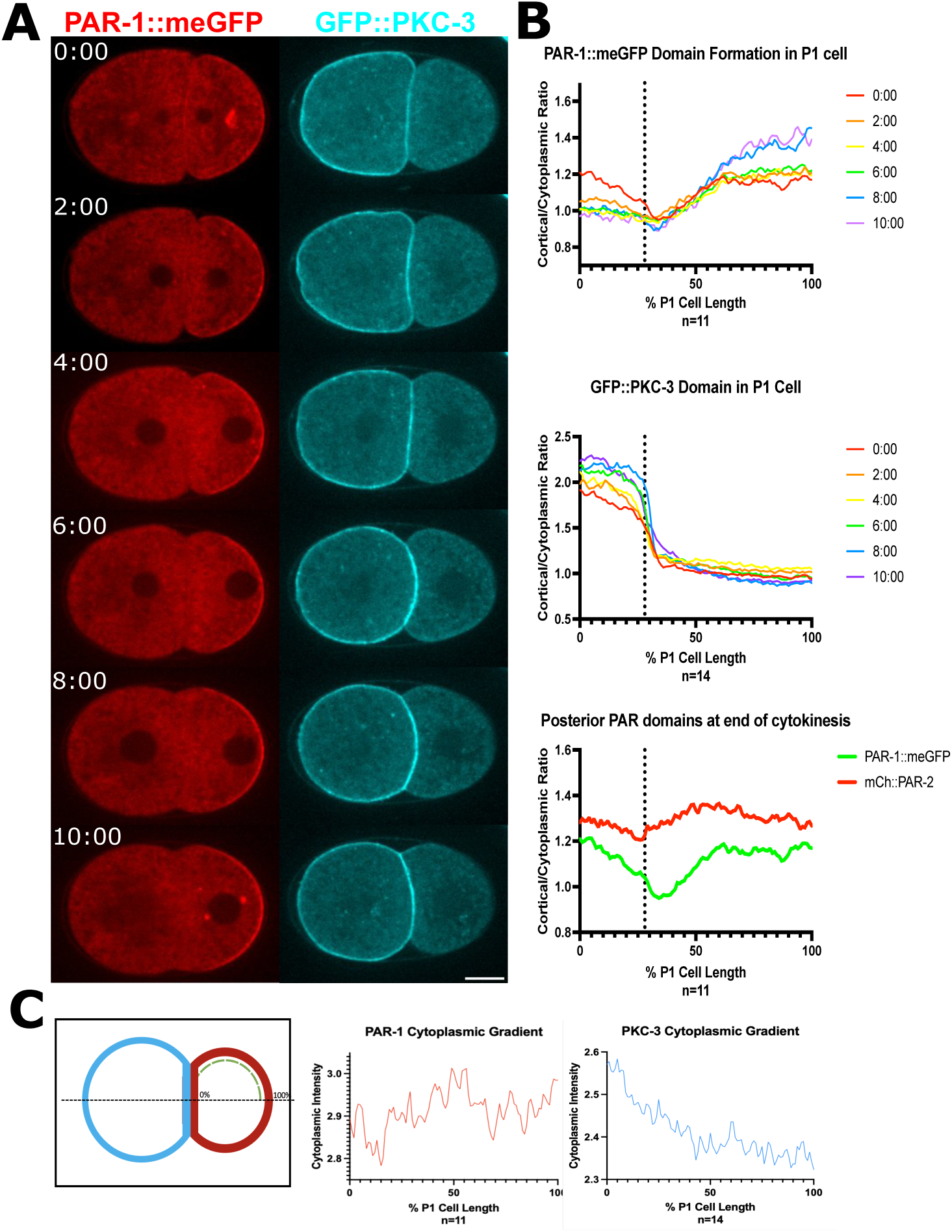
PAR-1 and PKC-3 are inherited asymmetrically in the P1 cell. (A) Confocal images of embryos expressing PAR-1::meGFP or GFP::PKC-3 . Time zero equals completion of cytokinesis. Scale bar is 10um. (B) Fluorescence intensity ratio plots measured as in Fig. 1, where each line represents a timepoint every two minutes for the first ten minutes; each line is an average of multiple embryos. Dotted line highlights the end of the AB-P1 cell contact. (C) Illustration of P1 cytoplasmic polarity measurements; embryos were divided along longest anterior-posterior axis and traced along green dotted line. (D) Average cytoplasmic intensity plots of PAR-1::meGFP and GFP::PKC-3 at 0 minutes after cytokinesis.

We also imaged embryos expressing GFP::PKC-3 and measured the cortical localization of PKC-3 in the P1 cell at the same time points (Fig 4A). Similar to PAR-6, PKC-3 appeared to only come into the furrow on the AB side (Fig S4A). PKC-3’s cortical levels increased globally in the first four minutes. Then at four minutes, PKC-3 started to clear from the posterior and continued to accumulate in the anterior (Fig 4B). We also looked at the cytoplasmic localizations of PAR-1 and PKC-3. We found that the P1 cell inherits a small gradient of PAR-1 in the posterior of the cell and a strong gradient of PKC-3 in the anterior (Fig. 4C). These data support a new hypothesis that the P1 cell inherits both cortical and cytoplasmic polarity from the P0 cell, and these inherited cues might be required for proper P1 cell polarization.

### *par-1* mutant embryos exhibit delayed polarization

Because of PAR-1’s distinct cortical localization pattern at the birth of the P1 cell, we hypothesized that PAR-1 has a role in P1 cell polarization. Previous work has shown that PAR-1 is redundantly required for cortical polarization in the P0 cell; in *par-1* mutant or RNAi embryos, aPAR and pPAR domains are normal at cytokinesis. This allowed us to examine the effect of loss of PAR-1 on P1 polarization. We first used RNAi of PAR-1 in the mCh::PAR-2 background. PAR-2 was inherited all around the P1 cell at cytokinesis completion, as in controls. However, in these embryos, clearing of PAR-2 from the anterior cell contact was substantially delayed and a PAR-2 domain was not apparent until 6-8 minutes after cytokinesis (Fig. 5A). Quantification of the posterior/anterior ratio of PAR-2 confirmed that there was no polarization of PAR-2 at 4 minutes and that polarization at 8 minutes was weaker than in controls (Fig. 5B). Analysis of PAR-6::GFP in *par-1 (RNAi)* embryos showed that there were higher levels of PAR-6 on the P1 cell cortex than in controls as the cell cycle proceeded, as predicted if PAR-1 inhibits cortical aPAR accumulation (Fig. S5). The delayed polarization of PAR-2 in the P1 cell was also exhibited by *par-1 (b274)* null mutant embryos. To test if PAR-1’s kinase activity is required for early clearing, we analyzed embryos with the kinase dead allele, *par-1 (it51)* (Guo & Kemphues, 1995). These embryos also exhibited a polarity delay (Fig. 5A-B). Together these results show that there is an “early” polarization pathway that depends on PAR-1 and its kinase activity and a “late” polarization pathway that doesn’t require PAR-1.

**Figure 5.**
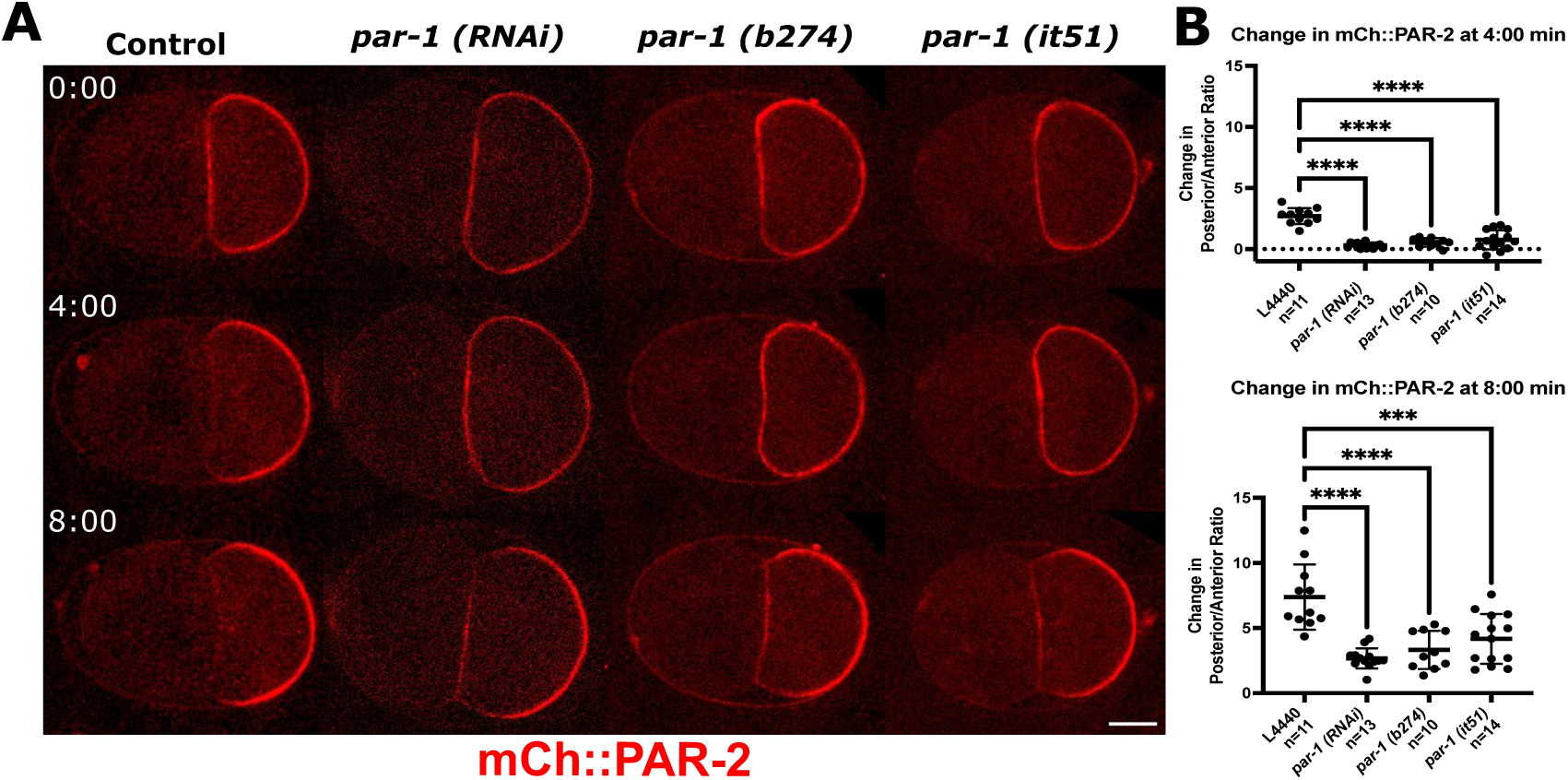
*par-1* mutants exhibit a polarization delay in the P1 cell. (A) Confocal fluorescent images of mCh::PAR-2 in L4440 control, *par-1 (RNAi), par-1(b274), and par-(it51)* embryos. Scale bar is 10um. See also Supplemental Videos 5 and 6. (B) Quantification of the change in posterior/anterior ratio of mCh::PAR-2 from 0 to 4 minutes and 0 to 8 minutes after cytokinesis. Means and statistics are reported in Supplemental Table 2.

Although *par-1* null mutants inherit normal cortical polarity at the two-cell stage, cytoplasmic polarity is inherited incorrectly (Griffin et al., 2011). To further examine the role of PAR-1 in P1 polarization, we examined existing PAR-1 mutants that have been reported to differentially affect cortical and cytoplasmic polarity: *par-1 (T983A), par-1 (KRSS),* and *par-1 (ΔKA1)* (Folkmann & Seydoux, 2019). In *par-1 (T983A)* mutants the PKC-3 phosphorylation site has been mutated; in *par-1 (KRSS)* and *par-1 (ΔKA1)* mutants, PAR-1’s autoinhibitory domain has been mutated at two important sites or deleted, respectively. In *par-1 (T983A)* embryos, PAR-1 is uniform on the cortex and at reduced levels; however cytoplasmic polarity is largely normal, likely due to normal inhibition of PAR-1 kinase activity at the anterior in this mutant. In *par-1 (KRSS)* and *par-1 (ΔKA1)* embryos, PAR-1 is no longer on the cortex in either mutant, but cytoplasmic polarity is more abnormal in *par-1 (ΔKA1)* compared to *par-1 (KRSS)* embryos (Folkmann & Seydoux, 2019). These *par-1* mutations were crossed into the mCh::PAR-2 background, and the cortical PAR-1 mislocalization phenotypes were verified in this new strain (Fig. S6). Interestingly, we found that the *par-1(ΔKA1)* mutant embryos showed the same mCh::PAR-2 polarization delay as *par-1* null mutants, while *par-1(T983A)* and *par-1 (KRSS)* embryos cleared PAR-2 similarly to controls at four minutes (Fig. 6A-B). These data support the hypothesis that the polarity delay in *par-1* mutants is the result of abnormal cytoplasmic polarity inherited from the P0 cell.

**Figure 6.**
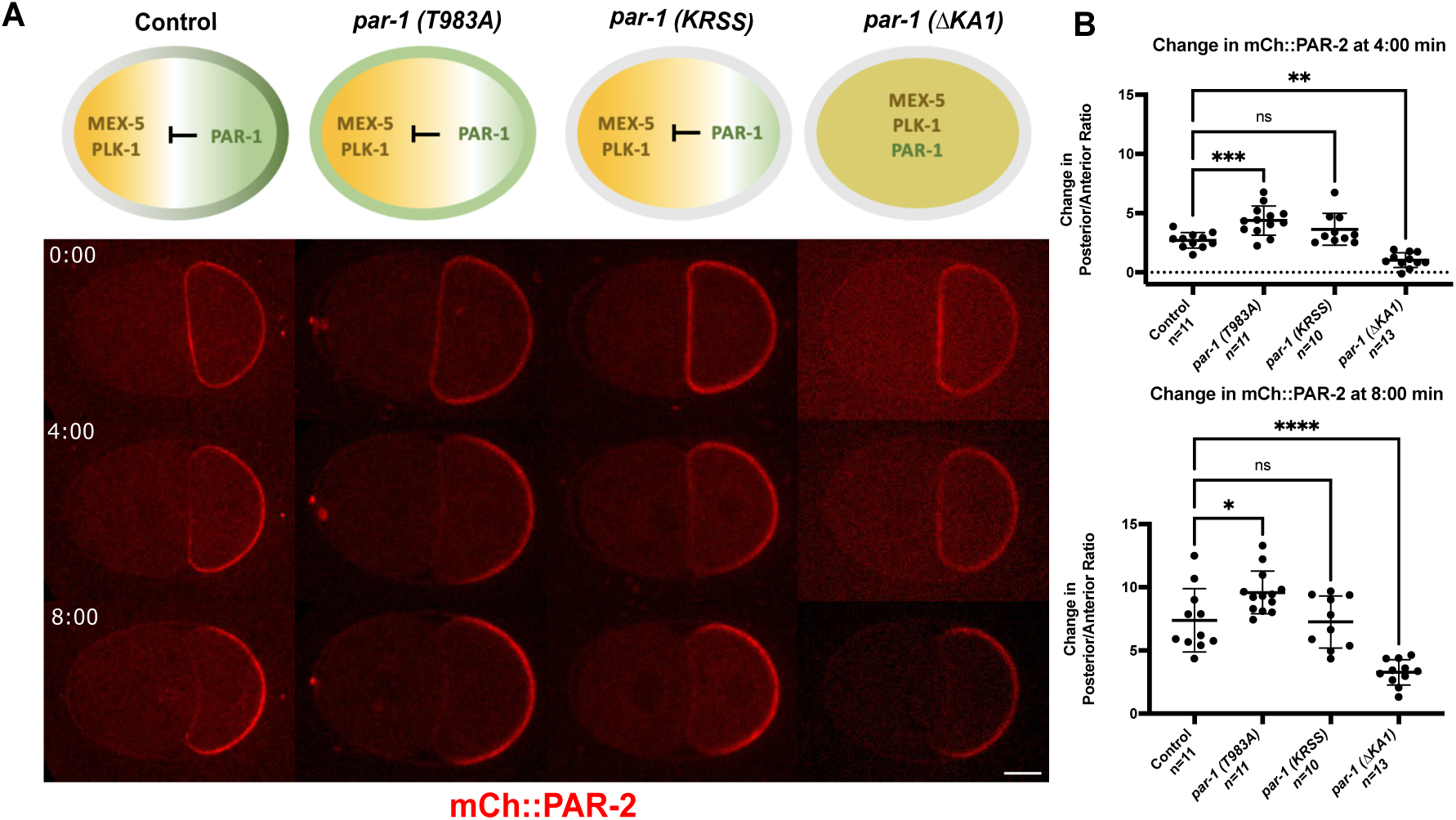
The delay in P1 polarization is caused by incorrect cytoplasmic polarity. (A) Diagrams of PAR-1, MEX-5, and PLK-1 distributions in controls and *par-1* mutants based on published studies. Confocal fluorescent images of mCh::PAR-2 in control, *par-1 (T983A), par-1(KRSS), and par-(ΔKA1)* embryos. Scale bar is 10um. (B) Quantification of the change in posterior/anterior ratio of mCh::PAR-2 from 0 to 4 minutes and 0 to 8 minutes after cytokinesis. Means and statistics are reported in Supplemental Table 2.

### *plk-1* mutant embryos exhibit delayed P1 polarization

To further test if the polarity delay seen in *par-1* embryos is the result of abnormal cytoplasmic polarity, we looked at downstream targets of PAR-1. MEX-5 and MEX-6 are highly similar zinc finger proteins that bind RNA, and they are required for generating the asymmetry of cell fate determinants downstream of PAR-1 (Schubert et al., 2000). PLK-1 is a mitotic kinase enriched in the anterior by MEX-5/6, and PLK-1 is also required for regulating the asymmetry of several cytoplasmic factors (Kim & Griffin, 2020). In wild-type one-cell embryos, PAR-1 has a posterior cytoplasmic gradient, while MEX-5/6 and PLK-1 localize in anterior cytoplasmic gradients; the AB cell inherits more MEX-5/6 and PLK-1 than the P1 cell (Cuenca et al., 2003; Guo & Kemphues, 1995; Nishi et al., 2008; Schubert et al., 2000) We verified that in our *par-1 (RNAi)* embryos, MEX-5 and PLK-1 were uniform in the one-cell embryo and that the AB and P1 cells received equal amounts (Fig. S7A-D). In *mex-5 (zu199); mex-6 (pk440)* mutant embryos, we found that the one-cell PAR-2 domain was smaller than in controls and PAR-2 did not flow into the furrow (Fig S7E). Because PAR-2 was not inherited uniformly in this background, we could not quantify polarization.

Next, we examined the role of PLK-1. PLK-1 was shown to inhibit cortical aPAR association in the oocyte and prevent premature symmetry breaking mutants (Reich et al., 2019). We therefor hypothesized that the high levels of PLK-1 in the P1 cell of *par-1* mutants might inhibit symmetry breaking and cause the late polarization in the *par-1* embryos. To test this hypothesis, we examined a *plk-1 (or683)* temperature sensitive mutant (O’Rourke et al., 2011) to determine if loss of PLK-1 could restore normal polarization kinetics in *par-1(RNAi)* embryos. Because PLK-1 also affects cell cycle timing, we quantified the change in polarization relative to P1 cell cycle length, where four minutes in controls equals 25% of the cell cycle, and eight minutes equals 50%. We found that *plk-1 (or683)* mutant embryos exhibited a polarity delay that was similar to that observed in *par-1(RNAi)* embryos (Fig. 7A, B). Further, the loss of PLK-1 neither rescued nor enhanced the polarity delay in combination with *par-1 (RNAi)*. These results suggest that excess PLK-1 does not cause the polarity delay in *par-1 (RNAi)* embryos. Rather, the data suggest that *par-1 (RNAi)* and *plk-1 (or683)* act in the same pathway to regulate P1 cell polarity and supports the model that normal one-cell cytoplasmic polarity is required for polarity of the P1.

**Figure 7.**
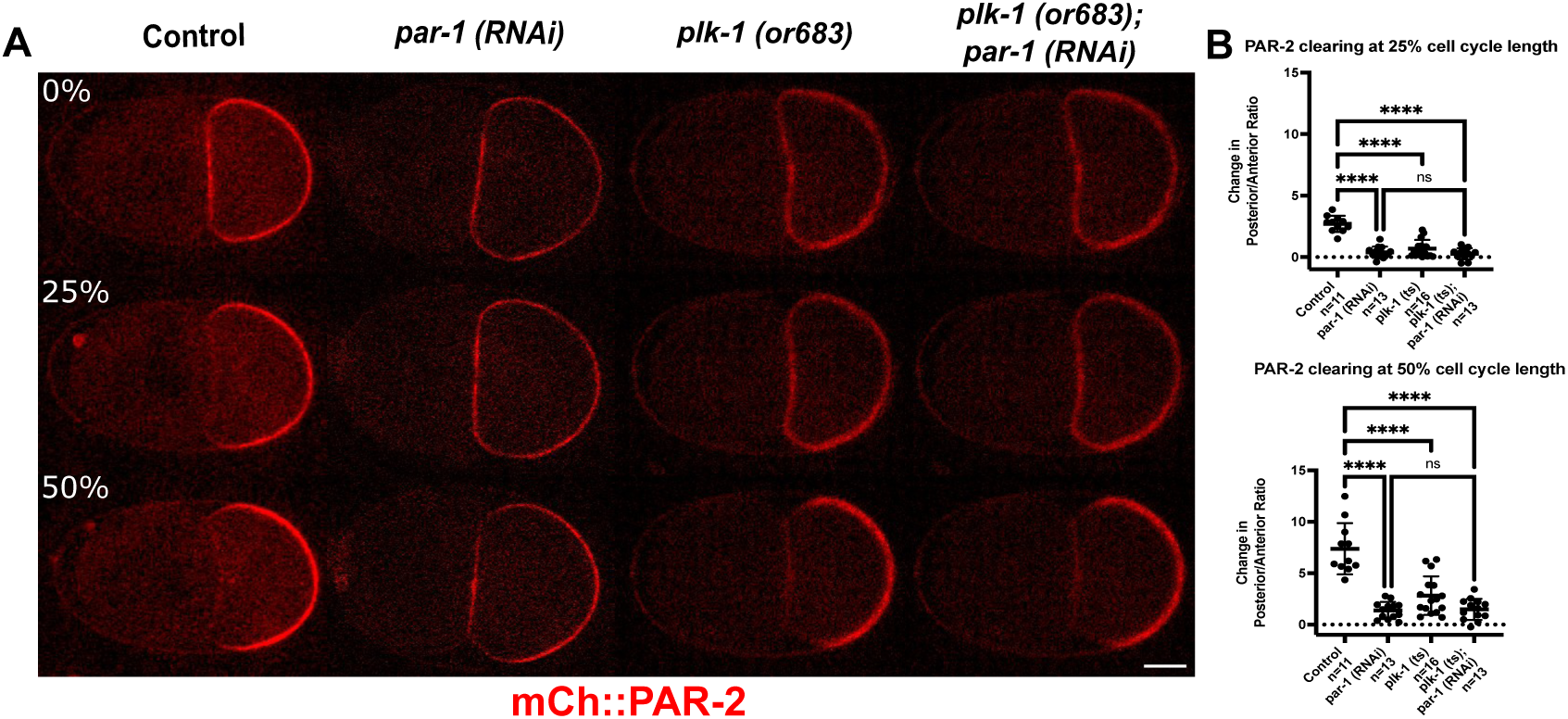
*plk-1* mutants exhibit a delay in P1 polarization. (A) Confocal fluorescent images of mCh::PAR-2 in control, *par-1 (RNAi), plk-1 (or683ts), plk-1 (or683ts); par-1 (RNAi)* embryos. *plk-1 (or683ts)* embryos were shifted to room temperature 30 minutes before filming. Scale bar is 10um. (B) Quantification of the change in posterior/anterior ratio of mCh::PAR-2 from 0%, 25%, and 50% P1 cell cycle length relative to total cell cycle length in each treatment. See Supplemental Table 1 for cell cycle lengths for each genotype. Means and statistics are reported in Supplemental Table 2.

### NMY-2 is required for late polarization

Although *par-1* mutant embryos exhibit a polarity delay, they do form a PAR-2 domain by eight minutes after cytokinesis. Based on our analysis of the timing of nuclear movement and actomyosin flow, we hypothesized that late polarization could be due to actomyosin flow. To test this, we carried out *par-1 (RNAi)* on the *nmy-2(ne3409ts) mCh::PAR-2* strain and shifted embryos to the restrictive temperature at the end of cytokinesis. The *par-1 (RNAi); nmy-2(ne3409ts)* embryos exhibited as stronger defect in P1 polarization at eight minutes, with a lower posterior/anterior ratio of PAR-2, than observed for *par-1 (RNAi)* alone (Fig. 8A-B). Further, 78.5% of *par-1 (RNAi); nmy-2(ne3409ts)* embryos also initiated PAR-2 clearing from the side instead of the AB/P1 cell contact (n=11/14) (Fig. S8). We also observed that while *nmy-2(ne3409ts)* single mutant embryos clear PAR-2 from the contact like controls at four minutes, at eight minutes they show significantly less polarization than controls. These results, together with the timing of myosin flow (Fig. 3) suggest that NMY-2 has a role in P1 cell polarization, but this pathway is not active until at least four minutes after cytokinesis.

**Figure 8.**
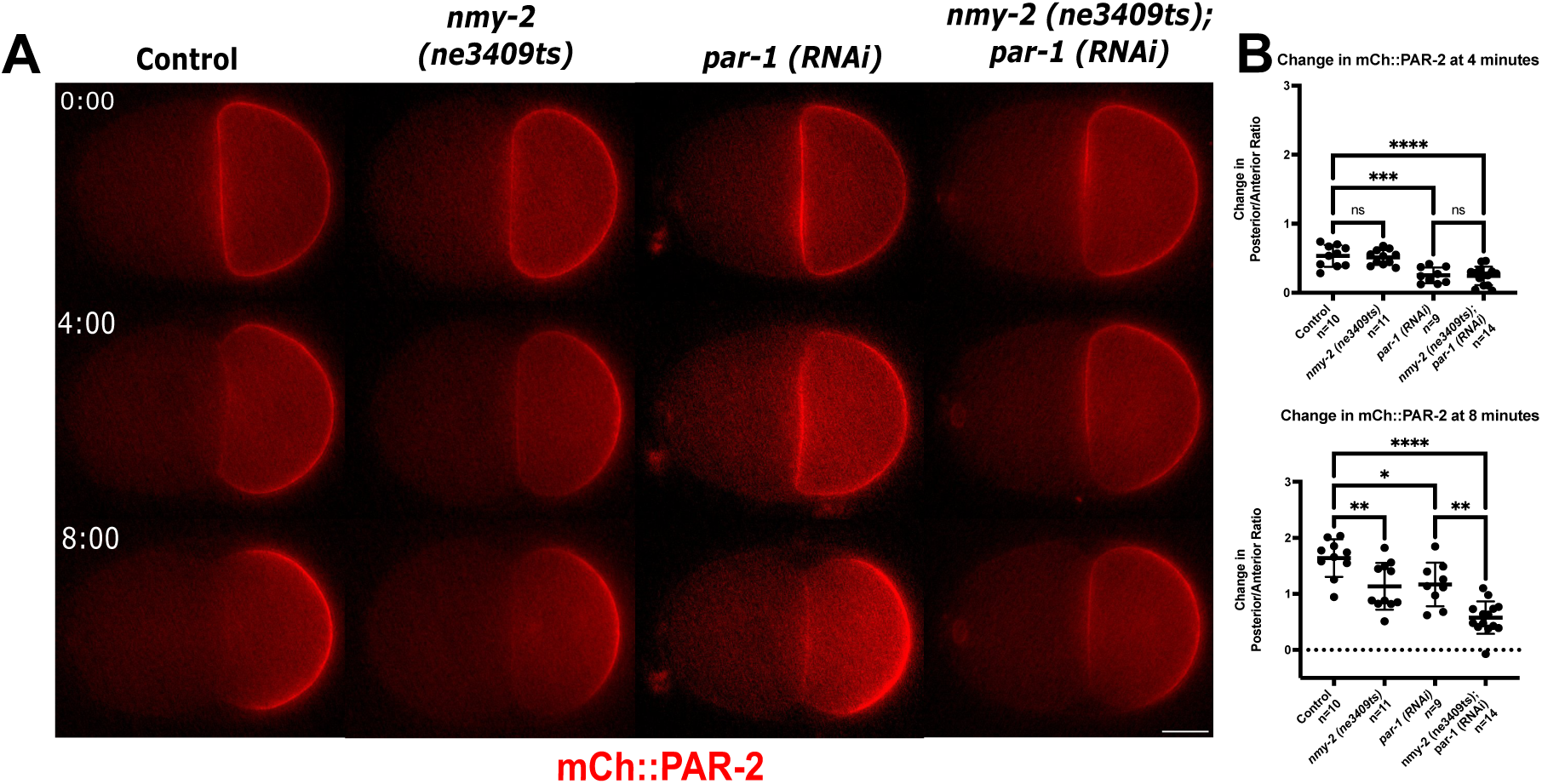
NMY-2 is required for late polarization in the P1 cell. (A) Epiflourescent images of mCh::PAR-2 in control, nmy-2*(ne3409), par-1 (RNAi),* and nmy-2*(ne3409); par-1 (RNAi)* embryos shifted from 16C to 26C at the end of P0 cytokinesis. Scale bar is 10um. (B) Quantification of the change in posterior/anterior ratio of mCh::PAR-2 from 0 to 4 minutes and 0 to 8 minutes after cytokinesis. Means and statistics are reported in Supplemental Table 2.

### Loss of PKC-3 blocks P1 cell polarization

To further test the mechanism by which PAR-2 is cleared from the P1 cell, we tested whether PKC-3 plays a role, using a temperature sensitive mutant, *pkc-3 (ne4250ts)* (Fievet et al., 2013), in the mCh::PAR-2 background. Even when grown and imaged at 16C, some *pkc-3 (ne4250ts);* mCh::PAR-2 embryos exhibited *pkc-3* mutant phenotypes such as a symmetric first cell division and incorrect division patterns at the two-cell stage, suggesting that this allele is a hypomorph even at permissive temperature. Nevertheless, in such *pkc-3 (ne4250ts);* mCh::PAR-2 embryos, the one-cell embryo still formed a PAR-2 domain and the P1 cell inherited uniform PAR-2. Surprisingly at the two-cell stage, a PAR-2 domain never formed in the P1 cell (Fig. 9A-B), but in all embryos a PAR-2 domain formed in the AB cell by the time of AB cell cytokinesis (n =7, Fig. S9). These results indicate that PKC-3 is required for early PAR-2 domain formation in the P1 cell, but this data does not distinguish between a role for PKC-3 in one-cell polarity or acting more directly in the P1 cell.

**Figure 9.**
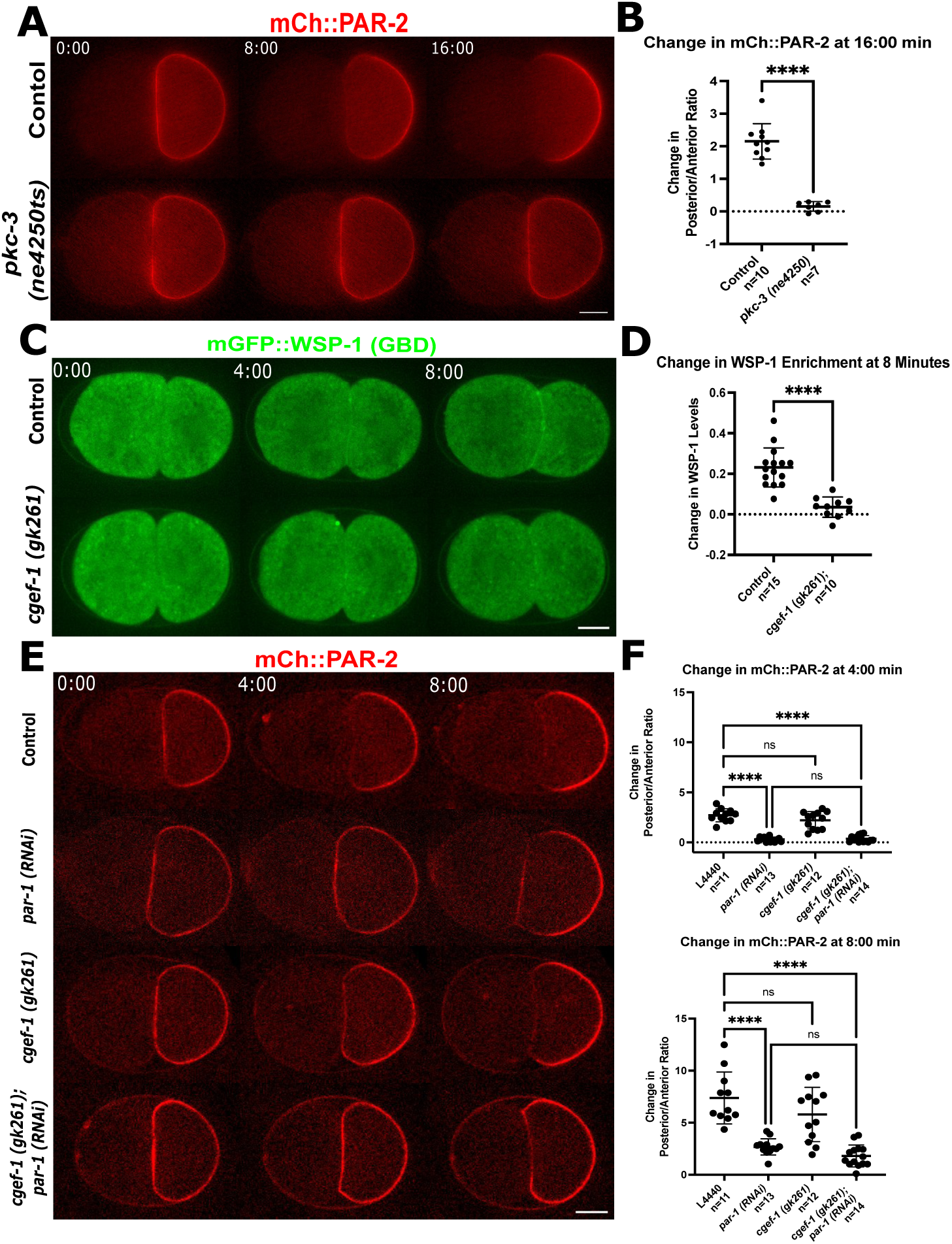
PKC-3 is required for early and late polarization in the P1 cell. (A) Epiflourescent images of mCh::PAR-2 in control and *pkc-3 (ne4250ts)* embryo grown and imaged at 16C. Scale bar is 10um. (B) Quantification of the change in posterior/anterior ratio of mCh::PAR-2 from 0 to 16 minutes after cytokinesis. (C) Confocal fluorescent images of mGFP::WSP-1GBD in control and *cgef-1 (gk261)* embryos. Scale bar is 10um. (D) Quantification of change in mGFP::WSP-1GBD at AB-P1 cell contract from 0 to 8 minutes relative to cytoplasm. (E) Confocal images of mCh::PAR-2 in control, *cgef-1(gk261), par-1 RNAi), and cgef-1(gk261); par-1(RNAi)* embryos. Scale bar is 10um. (F) Quantification of the change in posterior/anterior ratio of mCh::PAR-2 from 0 to 4 minutes and 0 to 8 minutes after cytokinesis. Means and statistics are reported in Supplemental Table 2.

We also tested whether another anterior PAR, CDC-42, contributes to PAR-2 clearing in the P1 cell. CDC-42 is a small GTPase that binds to PKC-3 and PAR-6 and is required for active PKC-3 in the one-cell embryo (Rodriguez et al., 2017; Seirin-Lee et al., 2020). To examine if CDC-42 is active in the P1 cell, we used a GFP tagged version of the WSP-1 G-protein-Binding-Domain, which is a published reporter for CDC-42 activity (Kumfer et al., 2010). In control embryos, active CDC-42 started to accumulate at the AB-P1 cell contact around 4 minutes after P0 cytokinesis and continued to accumulate throughout the cell cycle (Fig. 9C-D).

Because CDC-42 is required for proper P0 cell polarization we could not examine the P1 cell in a *cdc-42* null mutant. Instead, we examined embryos mutant for CGEF-1, a guanine nucleotide exchange factor that is partially redundant for activating CDC-42 in the early embryo (Kumfer et al., 2010). We first imaged GFP::WSP-1(GBD) in *cgef-1 (gk261)* null mutant embryos and found GFP::WSP-1(GBD) no longer accumulated on the AB-P1 cell contact (Fig. 9C-D). This result indicates that CGEF-1 activates CDC-42 in the P1 cell. We next examined mCh::PAR-2; *cgef-1(gk261)* embryos. These embryos exhibited partial *cdc-42* mutant phenotypes and were rounder than wild-type embryos, but the P0 cell had a normal PAR-2 domain at the posterior cortex (Kumfer et al., 2010). We observed that in *cgef-1(gk261)* embryos, P1 inherited PAR-2 uniform on the cortex, and polarization of the PAR-2 domain at four minutes was similar to controls. To test if CDC-42 has a role in the late pathway, we performed *par-1 (RNAi)* on mCh::PAR-2; *cgef-1(gk261)*. In this double mutant, we saw a more severe effect on polarization at eight minutes compared to *par-1 (RNAi)* alone (Fig. 9E-F). These results indicate that CDC-42, and by implication PKC-3, is required for the late polarization pathway.

## Discussion

The mechanisms by which polarity are established in the one-cell embryo (P0) of *C. elegans* have been extensively studied, as has polarization in other cell types in other organisms. However, much remains to be learned about how polarity is reestablished and maintained during successive asymmetric cell divisions and under different developmental conditions. Here, we characterized polarization of the P1 cell at the second division of the *C. elegans* embryo. In the P1 cell, reciprocal aPAR and pPAR domains form, but the cues for symmetry breaking and mechanism for polarization have not been investigated. We used mCh::PAR-2 as a marker for cortical polarity and confirmed that PAR-2 is inherited uniformly around the P1 cell. The polarization of P1 begins very early in the P1 cell cycle, with clearing of PAR-2 from the anterior AB/P1 cell contact region beginning within two minutes after P0 cytokinesis. Clearing continues and the levels of PAR-2 increase at the posterior so that a domain is visible at four minutes, but polarization of PAR-2 is stronger at eight minutes.

In the P0 cell, the presence of AIR-1 on the centrosome near the cortex appears to trigger symmetry breaking by inhibiting local myosin contractility; the resulting anterior directed myosin flow carries clusters of aPARs away from the centrosome and the presumptive posterior pole. PAR-1 and PAR-2 then associate with the posterior cortex (Klinkert et al., 2019; Munro et al., 2004; Schonegg et al., 2014; Zhao et al., 2019). Nuclear-centrosome movement towards the posterior and anterior-directed cortical myosin and PAR-6 flow have been observed in the P1 cell (Munro et al., 2004). However, here we found that strong cortical myosin flows occur well after the PAR-2 domain has started to form. Similarly, by analyzing AIR-1’s localization on the centrosome in P1, we conclude that AIR-1 is not in the correct position at the right time to act as a localized cue for early P1 polarization. Finally, using conditional mutants to inhibit AIR-1 recruitment to the centrosome or reduce myosin flow right after cytokinesis did not change the initial kinetics of PAR-2 clearing from the anterior cortex. These data together suggest that AIR-1 and actomyosin flow are not required for early polarization in the P1 cell.

Interestingly, we identified several asymmetries inherited by the P1 cell. Although PAR-2 is present uniformly around the P1 cortex after one-cell cytokinesis, the P1 cell is partially polarized for cortical PAR-1 and there are opposing cytoplasmic gradients of PAR-1 and PKC-3. It is also possible that low levels of PKC-3 are inherited in the P1 cell on the anterior cortex. However, due to the resolution limits of light microscopy, we cannot determine whether the signal on the AB/P1 contact is only in the AB cell or in both cells. Because PKC-3 and PAR-1 are known to inhibit each other’s localization, these inherited cytoplasmic or cortical gradients, where levels of PKC-3 are higher near the anterior, could trigger initial clearing of PAR-2. Consistent with this view, we found that *pkc-3(ne4250ts)* mutants showed normal cortical PAR-2 polarity at the end of P0 cytokinesis, but then never formed a normal PAR-2 domain. However, because the *pkc-3(ne4250ts)* allele has one-cell defects, these results are also consistent with a non-mutually exclusive model in which PKC-3 is needed for the proper asymmetry of other cytoplasmic components that play a role in P1 polarization, as outlined below.

Because of PAR-1’s initial asymmetry in the P1 cell, we also tested whether PAR-1 has a functional role in P1 cell polarization. Surprisingly the lack of PAR-1 cortical asymmetry or cortical localization did not affect P1 cell polarization. However, PAR-1’s kinase activity is required for early P1 cell polarization, and prior work showed that a PAR kinase activity gradient is required for normal cytoplasmic asymmetry (Folkmann & Seydoux, 2019; Griffin et al., 2011). Further, we found that *plk-1 (or683ts)* embryos and the *par-1 (RNAi); plk-1 (or683ts)* double mutants showed the same polarity delay as *par-1* mutant embryos. These observations lead us to the conclusion that PLK-1 and PAR-1 are acting in the same pathway to regulate a downstream cytoplasmic factor that is required for early polarization. We can envision two explanations for the polarity delay in *par-1* and *plk-1* mutants. One hypothesis is that there is a cytoplasmic gradient of a downstream target of PAR-1 and PLK-1 in the P1 cell, which acts as an inherited cue. and that cytoplasmic cue is lost in *par-1*and *plk-1* mutant embryos. Alternatively, there may be a cytoplasmic component that normally suppresses symmetry breaking in the AB cells. This component is mislocalized in the *par-1* and *plk-1* mutants such that high levels now suppress polarization of P1 as well.

Even though *par-1* mutant embryos do not polarize at the same time as wild-type embryos, they do eventually form a weak posterior PAR-2 domain. The time of AIR-1’s re-recruitment to centrosomes at the posterior and of NMY-2 flow in wild-type embryos correlates with the timing of late polarization observed in *par-1* mutants. Further, we found that simultaneous loss of PAR-1 and NMY-2 resulted in more severe polarization defects. In addition, *nmy-2* single mutants have a less robust PAR-2 domain at eight minutes. This leads us to propose that AIR-1 and NMY-2 have a role in the P1 cell, but they act in the late pathway, after the initial symmetry breaking event outlined above. It was previously shown that actomyosin flow corresponds with the movement of PAR-6, and our data is consistent with flow being the major driver of aPAR clearing in the posterior and accumulation in the anterior observed after four minutes wild-type embryo. In addition, the decrease in polarization of PAR-2 observed in *cgef-1* mutants, and the absence of late polarization in *pkc-3* mutants described, is consistent with PKC-3 playing a role in late polarization through exclusion of PAR-2.

All the data in this study supports a model in which there are two major pathways for timely polarization in the P1 cell (Fig. 10). There is a novel early pathway that initiates P1 polarization within two minutes after cytokinesis, which requires PAR-1, PKC-3, PLK-1, and the inheritance of normal cytoplasmic polarity. There is a second late pathway, which involves actomyosin flow-dependent aPAR accumulation in the anterior. This pathway enhances posterior PAR-1 and PAR-2 asymmetry in wild-type embryos as the cell cycle progresses and can function to polarize PAR-2 when early polarization is blocked. We also propose there are other pathways that can break symmetry in the P1 cell, because even in *par-1(RNAi); nmy-2(ts)* embryos, some PAR-2 clearing occurs late in the cell cycle. One possible mechanism for this clearing is that PAR-2 binding to microtubules emanating from the posterior centrosome at this time could protect it from phosphorylation by PKC-3; this pathway is redundant for P0 polarity (Motegi et al., 2011). In many *par-1(RNAi); nmy-2(ts)* embryos however, PAR-2 did not clear from the anterior AB-P1 cell contact as in controls, but instead cleared laterally, in the anterior corners. It has been previously reported that in the absence of the normal cue in the one-cell *C. elegans embryos*, there are other mechanisms that can spontaneously break symmetry that are influenced by cell shape (Klinkert et al., 2019); this phenomenon might be yet another way to break symmetry in the P1 cell.

**Figure 10.**
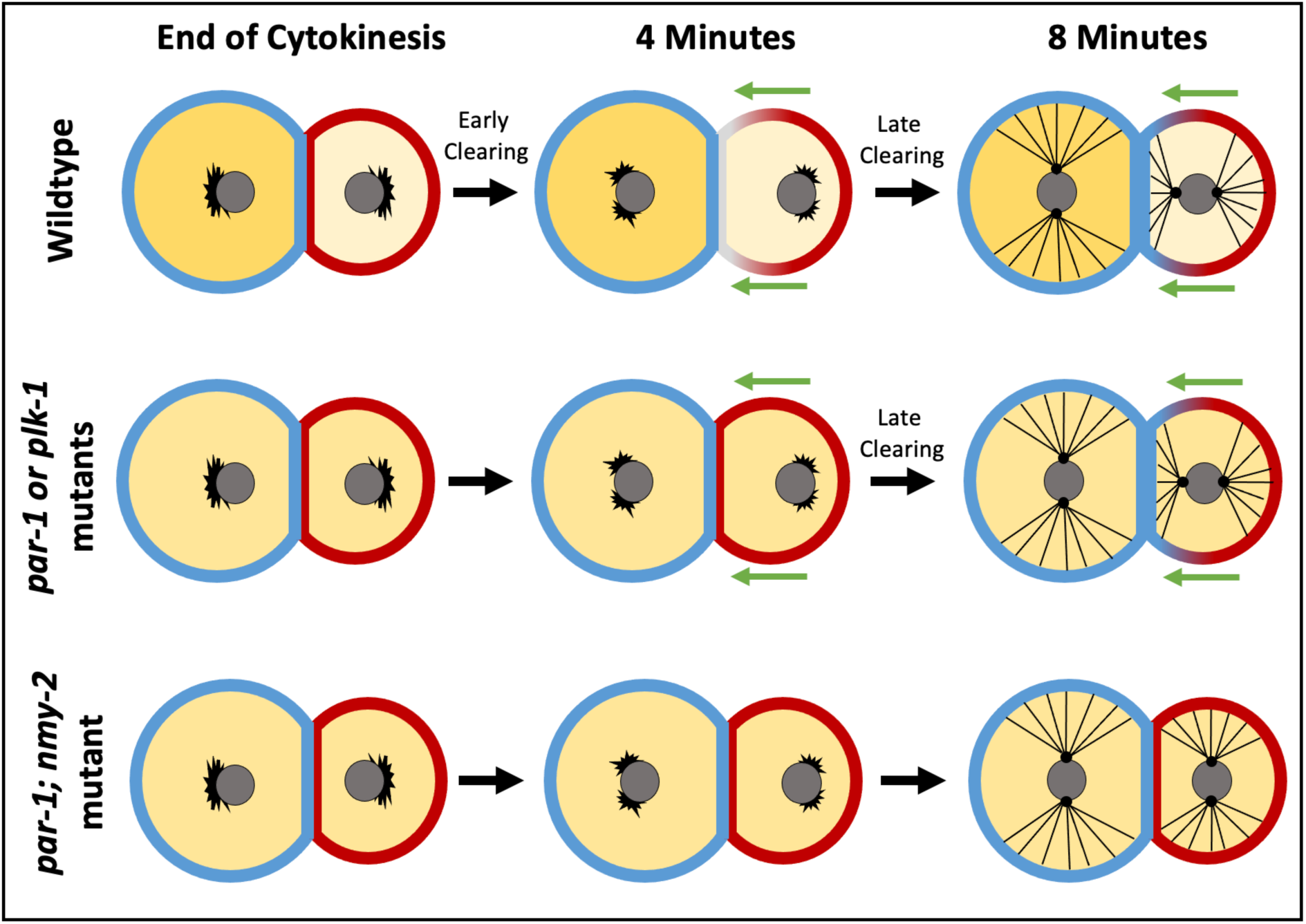
Model for Polarity Establishment in the P1 cell. Illustration of two pathways for establishing polarity at the 2-cell stage. Stages of polarization are shown at three timepoints in wildtype, *par-1* or *plk-1* mutants, and *par-1; nmy-2* mutants. PAR-2 is in red, PAR-6 is in blue, and cytoplasmic factors are in yellow; the grey circles are nuclei, the black lines shows microtubules emanating from the centrosome and the green arrows illustrate actomyosin flow.

In summary, our results identify a novel PAR-1 and PLK-1 dependent mechanism for polarization in the C. elegans embryo, which gives new insight into how cells in different developmental contexts can establish PAR polarity. The results also build on previous work in the C. elegans one-cell and Drosophila neuroblasts showing that cells employ multiple partially redundant pathways to promote robust polarization during asymmetric division.

## Materials and Methods

### *C. elegans* strains

*C. elegans* strains were maintained on MYOB plates with *E. coli* OP50 as a food source (Brenner, 1974; Church et al., 1995). The following strains were used in this study, listed in the order they appear in the paper:

| *Strain #* | *Genotype* | *Source* |
| --- | --- | --- |
| N2 | Wild type, Bristol Strain | CGC |
| KK1264 | *par-6(it310[par-6::gfp]) I; par-2(it315[mCherry::par-2]) III* | Ken Kemphues, (Reich et al., 2019) |
| RL439 | *par-2(it315) III; Itls78[(pKO5) pie-1::GFP::TEV::Stag::air-1 spliced coding + unc-119(+)]* | This paper |
| RL450 | *spd-5(or213ts) I; it315[mch::PAR-2] III; Itls78[(pKO5) pie-1::GFP::TEV::Stag::air-1 spliced coding* | This paper |
| JH2759 | *unc-119(ed3) III; axls1929[nmy-2::GFP + mCherry::par-2]* | (Zonies et al., 2010) |
| RL530 | *axls1929[nmy-2::GFP + mCherry::par-2]; par-2(it315[mCH::PAR-2]) III* | This paper |
| KK1254 | *par-2(it315[mCherry::par-2]) III* | Ken Kemphues |
| RL497 | *nmy-2 (ne3409ts) I; mch::par-2(it315) III* | This paper |
| JH3616 | *par-1(ax4206[PAR-1::meGFP]) V* | (Folkmann & Seydoux, 2019) |
| KK1228 | *pkc-3(it309 [GFP::PKC-3]) II* | Ken Kemphues |
| RL444 | *par-2(it315[mCherry::par-2]) III; par-1(ax4206[PAR-1::meGFP]) V* | This paper |
| RL520 | *par-2(it315[mCherry::par-2]) III; par-1 (it51) rol-4/ntl V* | This paper |
| RL521 | *par-2(it315[mCherry::par-2]) III; par-1 (b274) rol-4/ntl V* | This paper |
| RL542 | *par-2(it315[mCherry::par-2]) III; par-1(ax4207[PAR-1(K1170S R1171S)::meGFP])* | This paper |
| RL543 | *par-2(it315[mCherry::par-2]) III; par-1(ax4208[PAR-1(DKA1)::meGFP])/nT1[qIs51(pha::GFP)] V* | This paper |
| *RL544* | *par-2(it315[mCherry::par-2]) III; par-1(ax4209[PAR-1(T983A)::meGFP])/nT1[qIs51(pha::GFP)] V* | This paper |
| *RL545* | *par-2(it315[mCherry::par-2]) III; plk-1 (or683ts) III* | This paper |
| *RL473* | *pkc-3 (ne4250ts) II; par-2(it315[mCherry::par-2]) III* | This paper |
| *WH517* | *ojIs40 [wsp-1(GBD)::GFP + unc-119(+)]* | (Kumfer et al., 2010) |
| *WH527* | *cgef-1(gk261) X; ojIs40 [wsp-1(GBD)::GFP + unc-119(+)]* | (Kumfer et al., 2010) |
| *RL533* | *par-2(it315[mCH::PAR-2]) III; cgef-1 (gk261) X* | This paper |

### Live Imaging

Because all of the proteins under study are maternally provided, embryos were derived from homozygous mutant hermaphrodites, or hermaphrodites treated for RNAi, in all cases. Embryos were removed from gravid hermaphrodites, dissected into egg buffer (25 mM HEPES, pH 7.4, 120 mM NaCl, 48 mM KCl, 2 mM CaCl2, MgCl2), mounted on 2% agar pads, and covered with coverslip.

Epifluorescent microscopy was carried out on an Olympus BX60 microscope equipped with PlanApo N 60X, 1.42 NA oil immersion objective lens, a CoolLED light source, a Hammatasu Orca 12-bit digital camera, and MicroManager software. All time-lapse videos were taken at 10 seconds intervals, except for temperature sensitive mutants and their controls which were taken every 30 seconds.

Confocal microscopy was carried out using the spinning disc module of an Intelligent Imaging Innovations (3i) Marianas SDC Real-Time 3D Confocal-TIRF microscope fit with a Yokogawa spinning disc head, a 60x 1.4 numerical aperture oil-immersion objective, EMCCD camera, and Slidebook 6 software. Images were taken in a mid-focal plane at 10 seconds intervals, except for cortical images of NMY-2 for which 3 Z-planes were imaged with 0.5-micron steps every 3 seconds.

### RNAi and Temperature Sensitive Mutants

RNAi was performed by feeding (Timmons & Fire, 1998). The *par-1 (RNAi)* construct used was obtained from the Ahringer RNAi library (Kamath et al., 2003). RNAi was conducted for 48hrs at 20C to obtained published strong loss of function phenotypes, such as synchronous two-cell divisions and symmetric P0 cell division.

Fast-inactivating temperature sensitive mutants were filmed on a temperature control stage. Embryos were grown and dissected at 16°C prior to shifts. Temperature shifts were done by fitting a Linkam PE95/T95 System Controller with an Eheim Water Circulation Pump set and maintain the temperature of the slide. Stage was set to 12°C for 16°C and 26°C for 26°C, true temperatures were determined by inserting the wire probe of an Omega HH81 digital thermometer between the cover slip and an agar pad with oil on the 60X objective. The temperature control stage can shift the slide from 16C to 26C in 1 minute. The strength of each temperature sensitive mutant was compared to published mutant or loss of function phenotypes.

### Quantification

Analysis of mCh::PAR-2, PAR-6::GFP, PAR-1::meGFP, and GFP::PKC-3 domains in the P1 cell were done by dividing the embryo along its longest anterior-posterior axis. Then both the top and bottom cortices were traced using the segmented line tool (width = 2 pixels) in Fiji (as in Fig. 1B). Cytoplasmic mean was measured by drawing a small circle in they cytoplasm, avoiding the cortex and nucleus. Cortical intensity was divided by cytoplasmic mean and adjusted to percent P1 cell length. This was done for each embryo and then averaged to get a plot for each time point.

Analysis of cytoplasmic polarity in PAR-1::meGFP and GFP::PKC-3 was measured at the end of cytokinesis. As above, embryos were divided along their longest anterior-posterior axis, but then traces of a region just beneath the cortex were made using the segment line tool (width = 2 pixels) in Fiji, starting near the corner of the cell contact towards the most posterior point (as in Fig. 4C). Cytoplasmic intensity was divided by the background outside of the cell and adjusted to percent P1 cell length.

To analyze how close the centrosome moves towards to the posterior cortex, the frame where cytokinesis ended and then the frame in which the nucleus or GFP::AIR-1 foci were closest to the membrane were scored. At this timepoint the distance from the edge of the P1 nucleus or the GFP::AIR-1 foci to the posterior membrane was measured. The distance of the nucleus or GFP::AIR-1 foci was normalized to P1 cell length (longest anterior-posterior axis) to account for differences in embryo size. Fiji line tool was used for all these measurements and data was analyzed in GraphPrism.

Analysis of the change in mCh::PAR-2 domain over time was quantified using the Fiji line segment tool (width = 2 pixels) to trace the AB-P1 cell contract and the same length of the posterior cortex at zero, four and eight minutes after cytokinesis. For embryos filmed on the 3i confocal microscope, cortical traces were normalized to cytoplasmic mean. Because of the large amount of out of focus fluorescence within the cell, for embryos filmed with epifluorescence the cortical traces were normalized to the background outside of the cell. The normalized posterior value was divided by the normalized anterior value to give an posterior/anterior ratio for each time point. The difference between time points was found (example: P/A ratio at 4:00 – P/A ratio at 0:00) to calculate the change over time. All data was analyzed in GraphPrism.

Change in GFP::WSP-1(GBD) was measured by tracing the AB-P1 cell contact using Fiji line segment tool (width = 2 pixels). This was normalized to cytoplasmic mean to get a normalized intensity for each time point. Then the normalized intensity at eight minutes was subtracted by the normalized intensity at 0 minutes to find the changed in GFP::WSP-1(GBD). All data was analyzed in GraphPrism.

The mCh::MEX-5 and PLK-1::GFP cytoplasmic ratios were determined by drawing a small circle in the AB cell and P1 cell, avoiding membranes and the nucleus. The average mean of the AB cytoplasm was divided by the P1 cell to find the AB/P1 ratio. All data was analyzed in GraphPrism.

## Acknowledgements

We thank Christy Kok for help with filming N2 embryos and data analysis, Aidan van Cleef and Kathie Urrutia-Paniagua for media preparation, and members of the Rose and McNally labs for helpful discussions. We are grateful to Ken Kemphues, Geraldine Seydoux for providing strains. Other strains were provided by *Caenorhabditis* Genetics Center, which is funded by the National Institutes of Health Office of Research Infrastructure Programs (P40 OD010440). The 3i Marianas spinning disk confocal used in this study was purchased using a National Institutes of Health Shared Instrumentation Grant [1S10RR024543-01]. We thank the MCB Light Microscopy Imaging Facility, which is a UC-Davis Campus Core Research Facility, for maintaining this microscope. This research was funded by awards to LR from the National Institutes of Health Grant [R01GM68744] and the National Institute of Food and Agriculture [CA-D*-MCB-6239-H].

---

**Figure S1.**
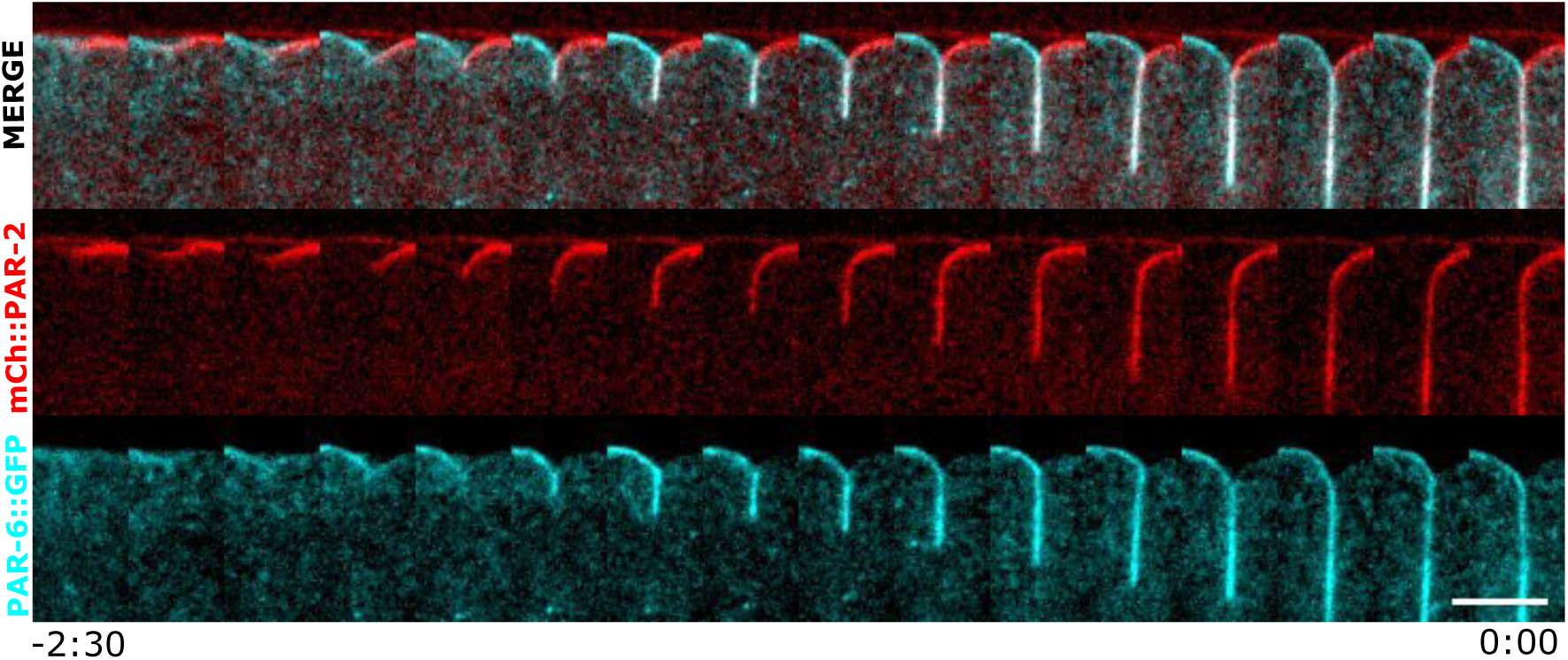
PARs during furrow ingression. Confocal images of the P0 cell cytokinetic furrow with mCh::PAR-2 and PAR-6::GFP. Each frame represents a 10 second interval. Anterior is to the left and posterior to the right in this and all supplemental figures. Scale bar is 5um.

**Figure S2.**
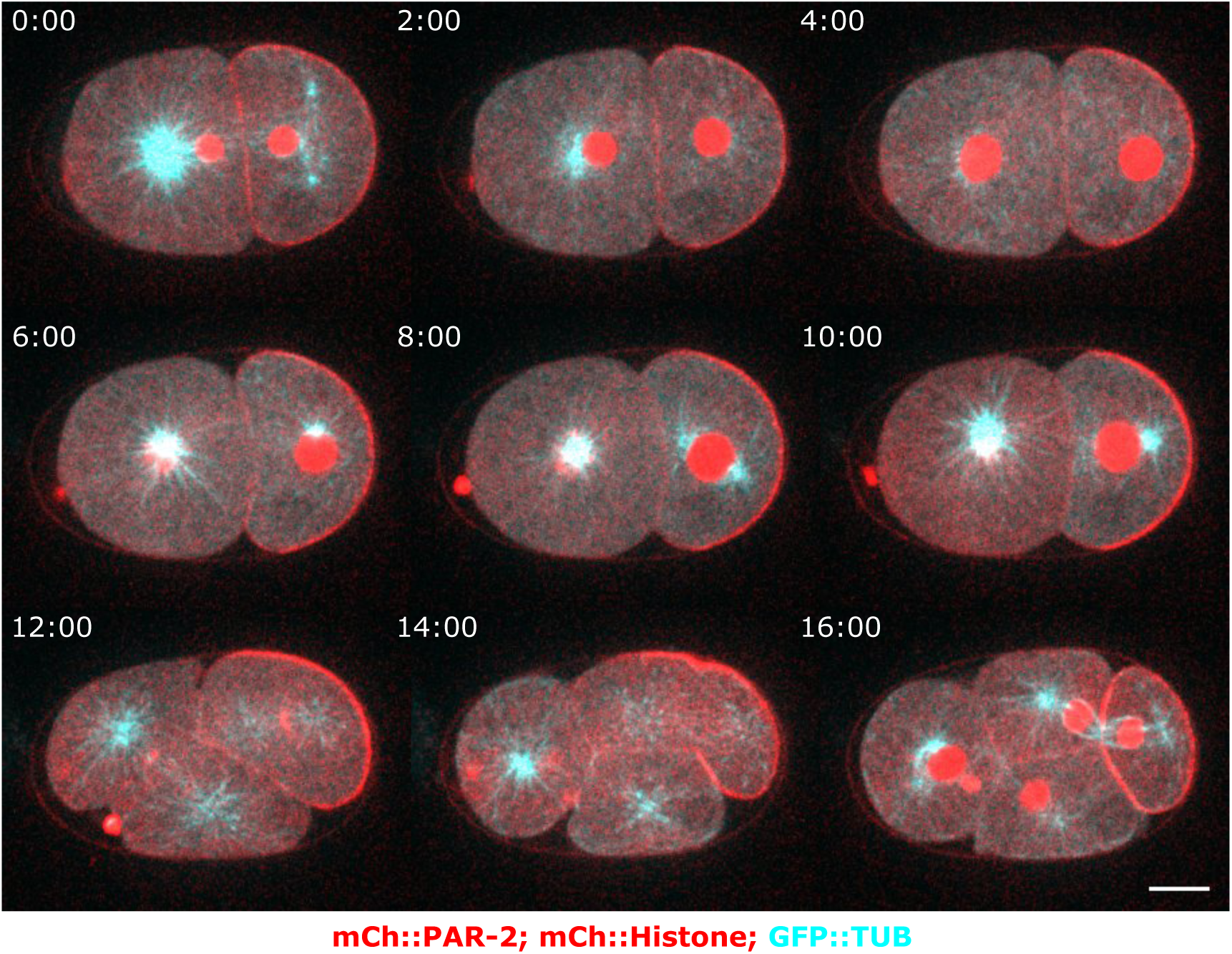
P1 cell cycle timing. Confocal images of embryos expressing mCh::PAR-2, mCh::Histone, and GFP::Tubulin labeled to illustrate cell cycle timing compared to P1 polarization. Each frame represents a 2 minute interval. Scale bar is 10um.

**Figure S3.**
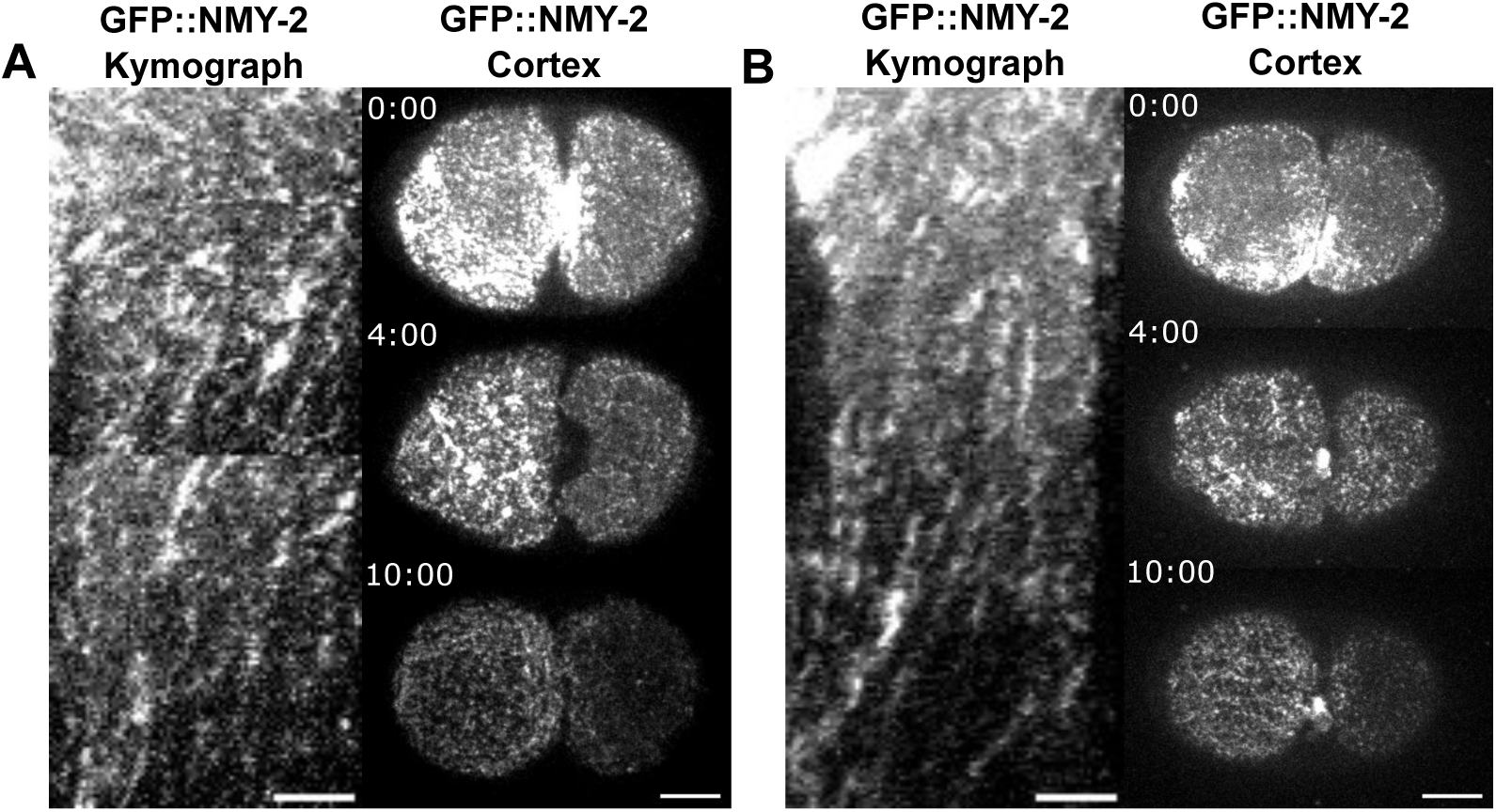
Actomyosin flow in P1 cell. (A-B) Additional examples of embryos expressing GFP::NMY-2. Kymograph of GFP::NMY-2 taken across the longest anterior-posterior axis. Confocal fluorescent images of the same embryo from a surface view at 0, 4, and 10 minutes after cytokinesis. Scale bar in kymograph is 5um and embryo is 10um.

**Figure S4.**
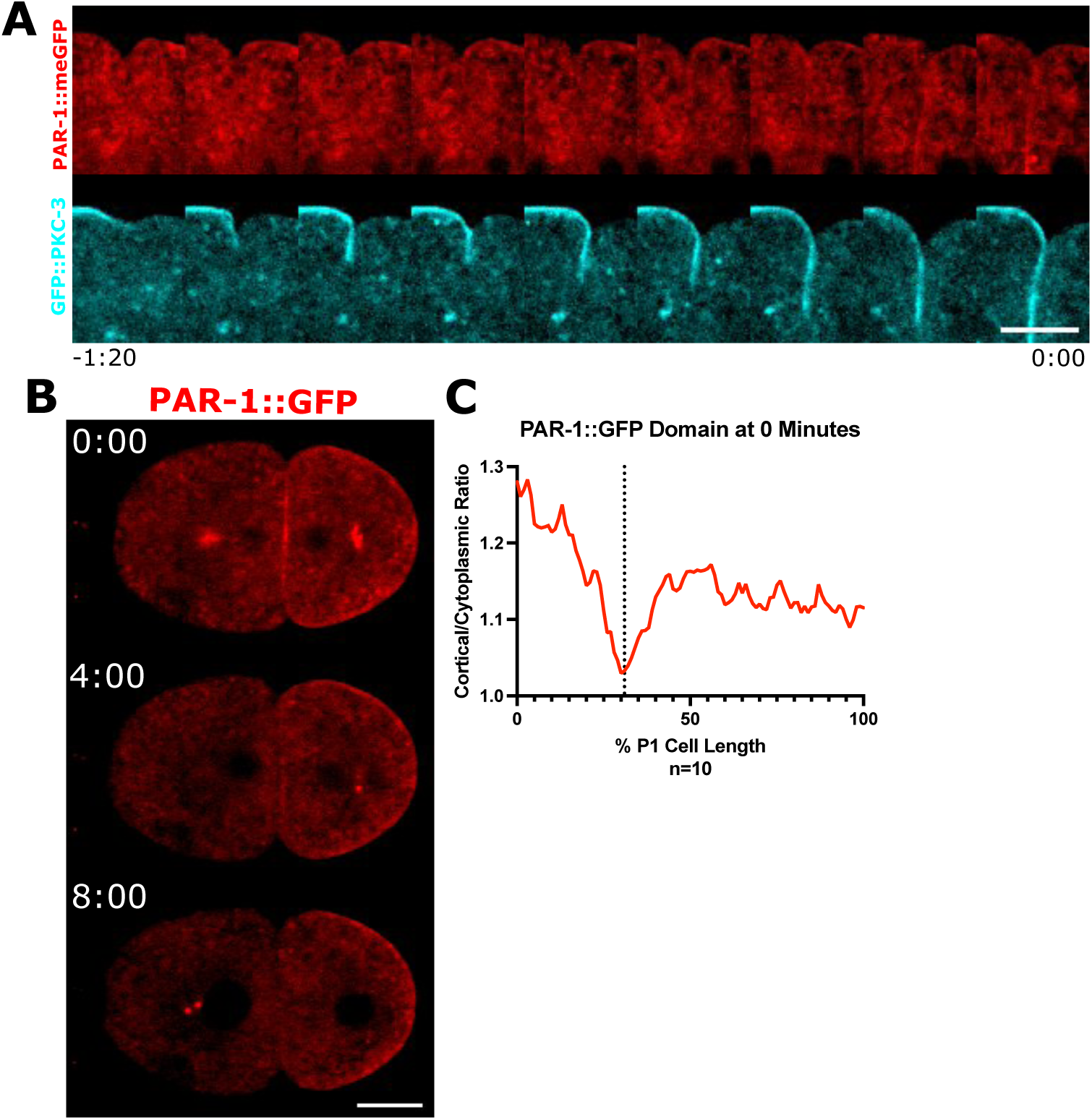
PAR-1 and PKC-3 during cytokinesis. (A) Confocal images of the P0 cell cytokinetic furrow with PAR-1::meGFP or GFP::PKC-3. Each frame represents 10 second interval. Scale bar is 5um. (B) Confocal images of embryos expressing a PAR-1::GFP transgene. Time zero equals completion of cytokinesis. Scale bar is 10um. (C) Quantifications of fluorescence intensity at the end of cytokinesis. Dotted line highlights the end of the AB-P1 cell contact.

**Figure S5.**
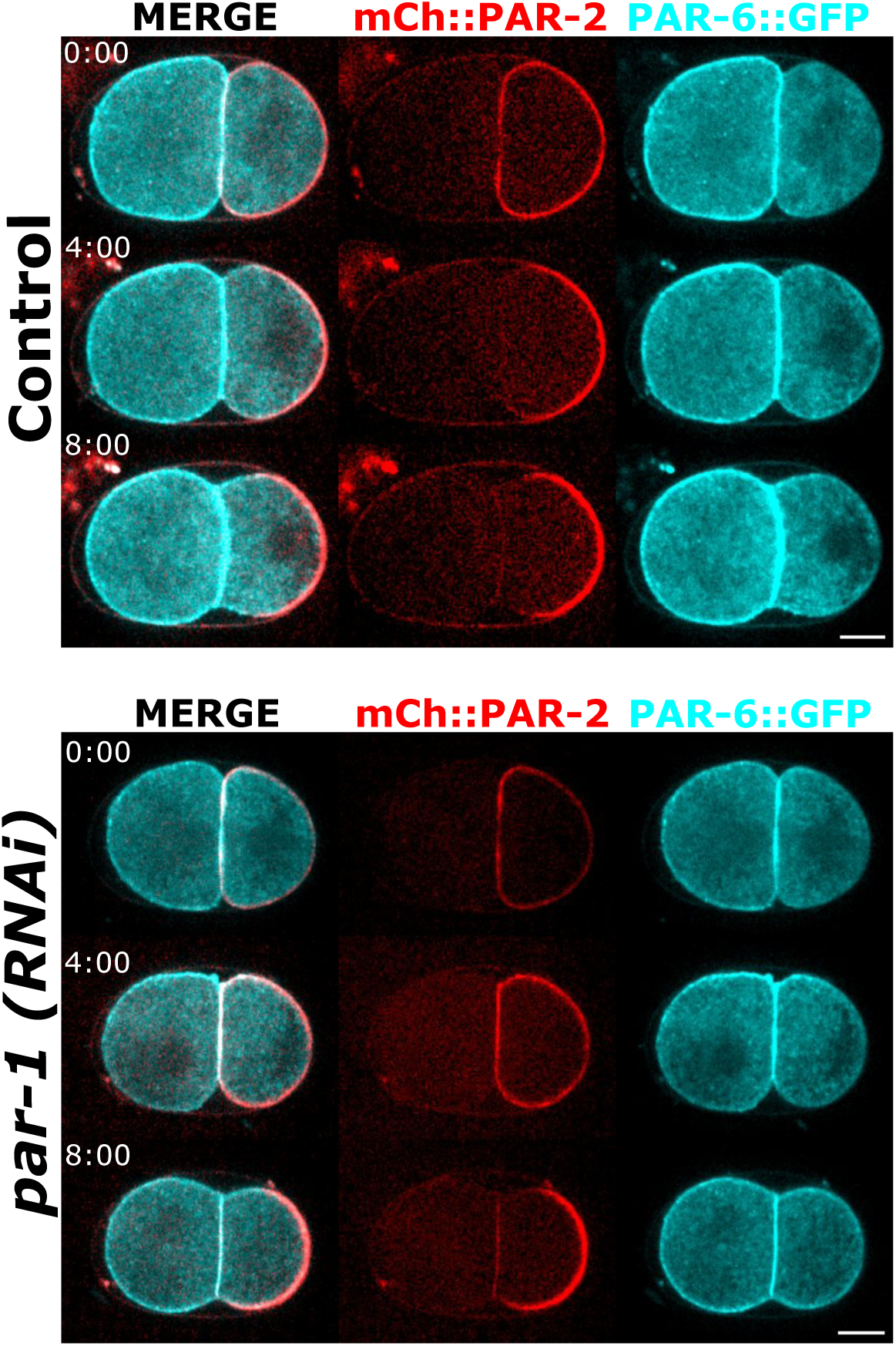
PAR-2 and PAR-6 in *par-1(RNAi) embryos*. Confocal fluorescent images of mCh::PAR-2 and PAR-6::GFP in L4440 control and *par-1 (RNAi).* Scale bar is 10um.

**Figure S6.**
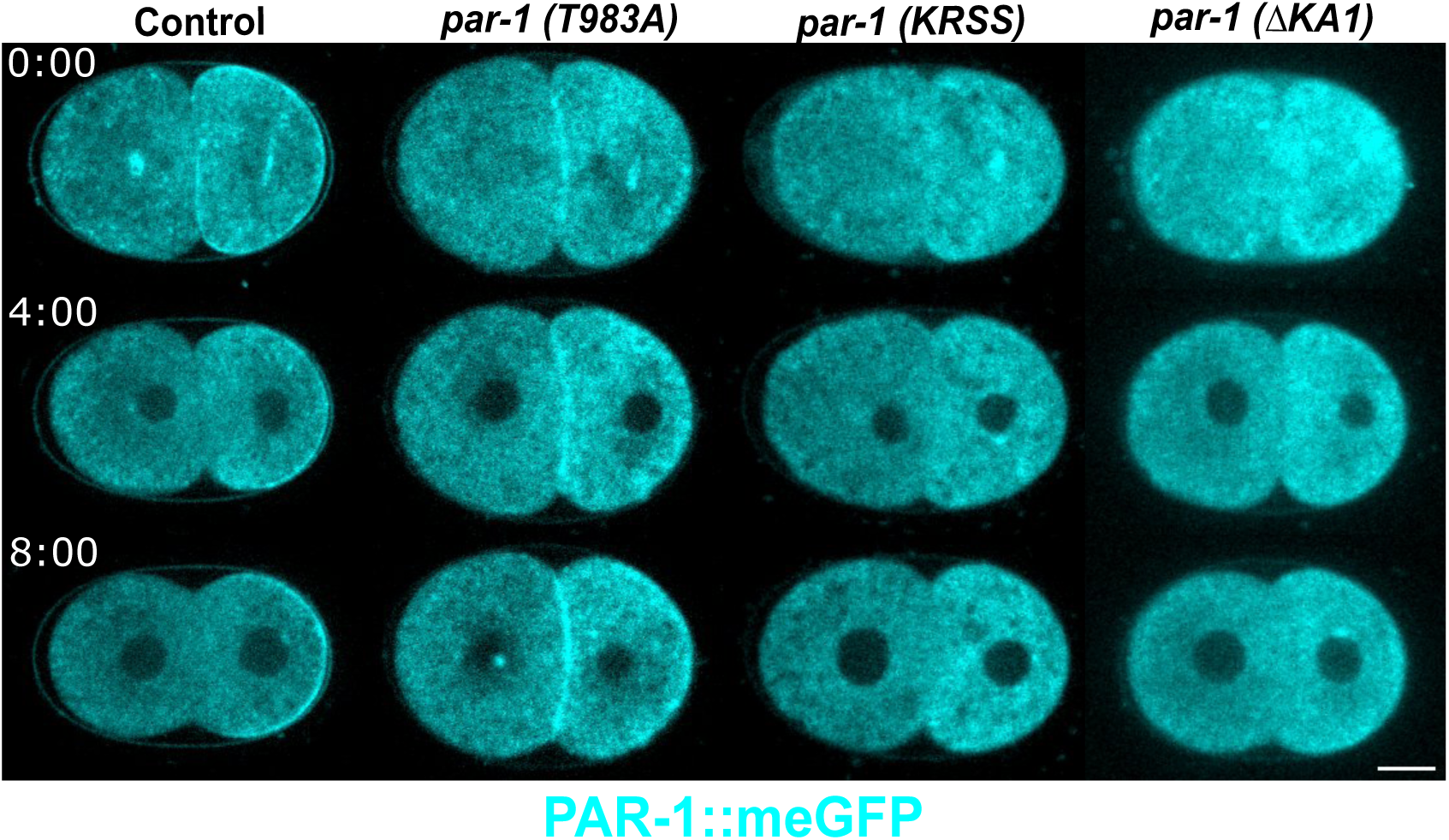
PAR-1 localization in *par-1* mutants. Confocal fluorescent images of PAR-1::meGFP, *par-1 (T983A)*::meGFP*, par-1(KRSS)*::meGFP*, and par-(ΔKA1)*::meGFP embryos at 0, 4, and 10 minutes after cytokinesis. Scale bar is 10um.

**Figures S7.**
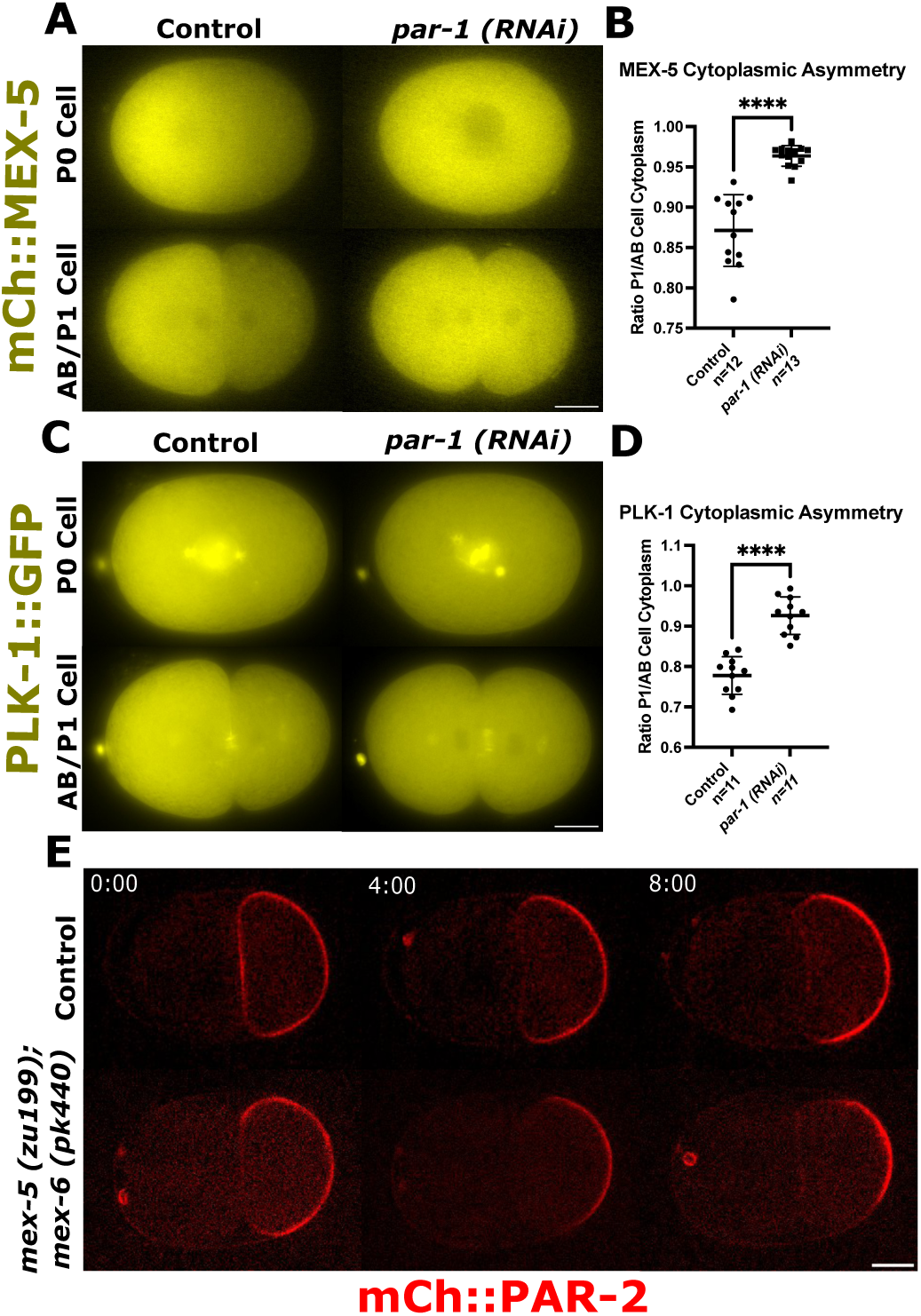
MEX-5 and PLK-1 cytoplasmic asymmetry at the two-stage is disrupted in par-1(RNAi) embryos. (A) Epiflourescent images of mCh::MEX-5 at the one-cell stage and at the end of cytokinesis. Scale bar is 10um. (B) Quantification of MEX-5 cytoplasmic asymmetry between the P1 and AB cells. (C) Epiflourescent images of PLK-1::GFP at the one-cell stage and at the end of cytokinesis. Scale bar is 10um. (D) Quantification of PLK-1 cytoplasmic asymmetry between the P1 and AB. (E) Confocal images of mCh::PAR-2 in control and *mex-5 (zu199); mex-6 (pk440)* embryos. Scale bar is 10um. Means and statistics are reported in Supplemental Table 1.

**Figure S8.**
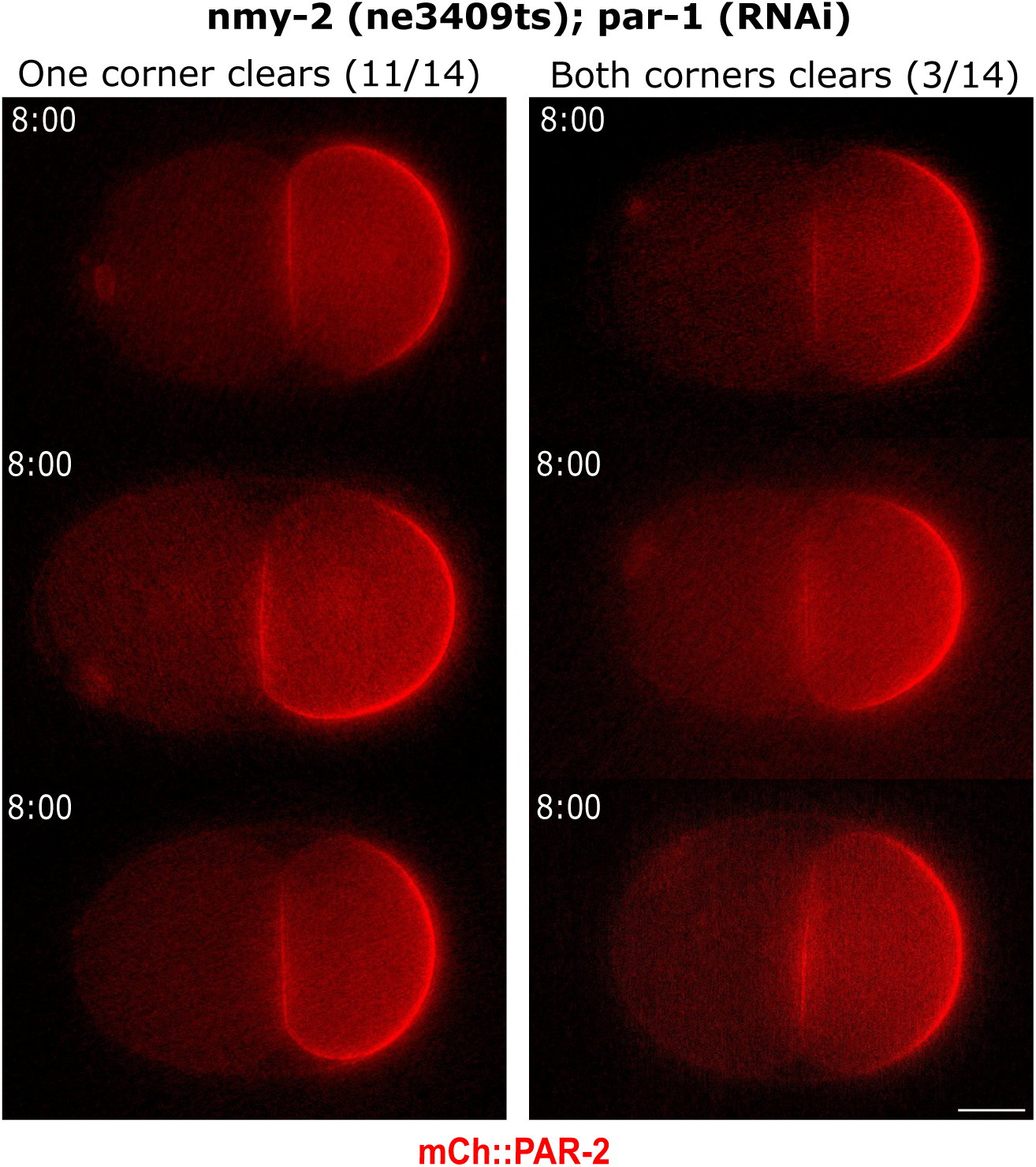
Different Patterns of PAR-2 Clearing in *nmy-2 (ne3409ts); par-1 (RNAi)* embryos. Representative images of *nmy-2 (ne3409ts); par-1 (RNAi)* embryos with mCh::PAR-2 at 8 minutes after cytokinesis. 11/14 embryos cleared from one corner and 3/14 embryos cleared in both corners. Scale bar is 10um.

**Figure S9.**
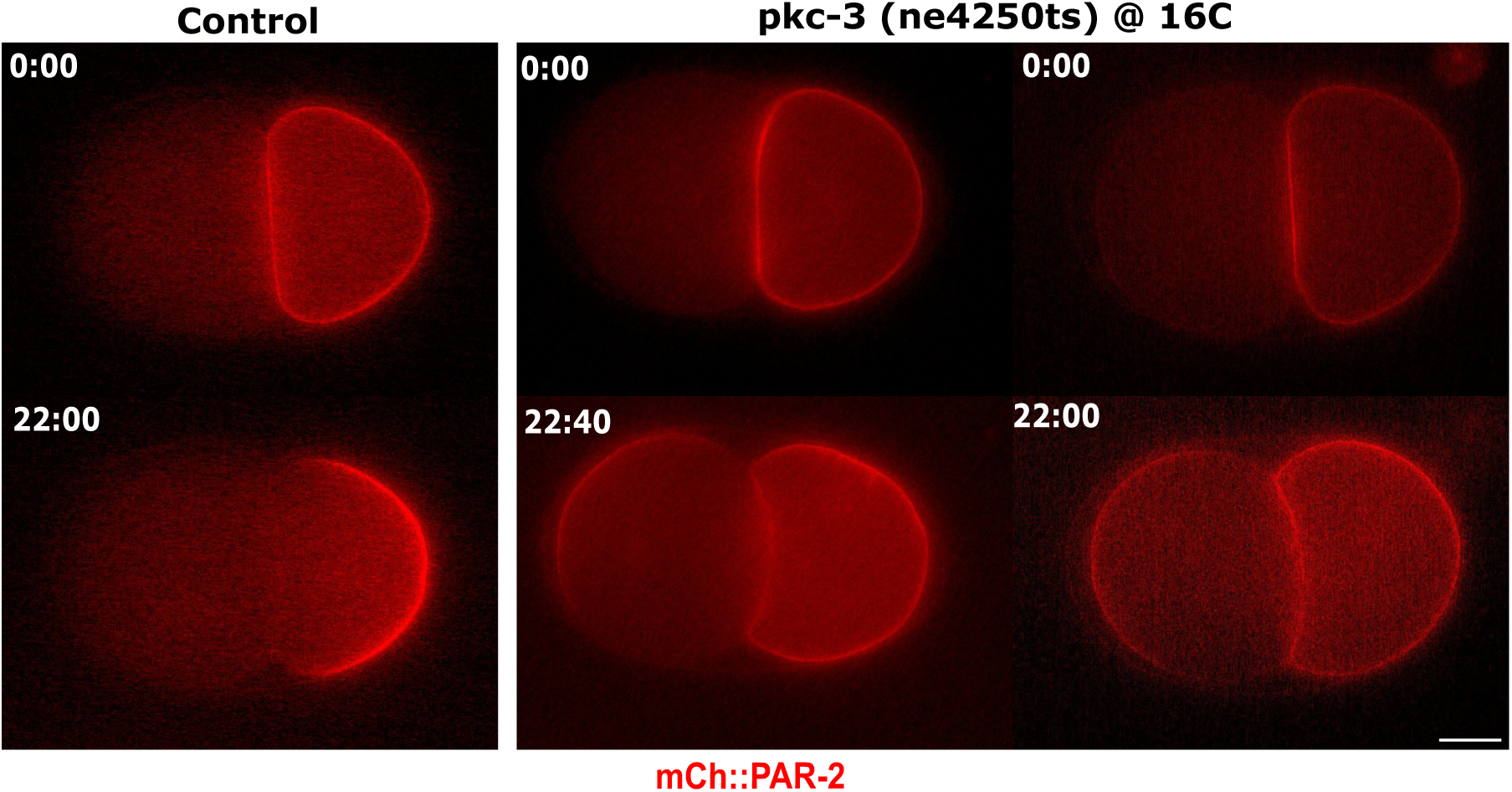
Ectopic PAR-2 domains form in the AB cell in *pkc-3 (ne4250ts).* Representative embryos expressing mCh::PAR-2 in WT or *pkc-3 (ne4250ts)* backgrounds filmed at 16C. Scale bar is 10um.

**Supplemental Table 1.** **–** Average cell cycle length in the different conditions.

| Supplemental Table 1 |  |  |  |
| --- | --- | --- | --- |
| Condition | Mean (minutes) | SD | n |
| mCh::PAR-2; Control | 16.07 | 1.395 | 11 |
| mCh::PAR-2; *par-1* (RNAi) | 13.31 | 1.562 | 7 |
| mCh::PAR-2; *plk-1* (*or683ts*) | 20.82 | 1.897 | 13 |
| mCh::PAR-2; *plk-1* (*or683ts*); *par-1* (RNAi) | 17.41 | 0.659 | 9 |

**Supplemental Table 2.**
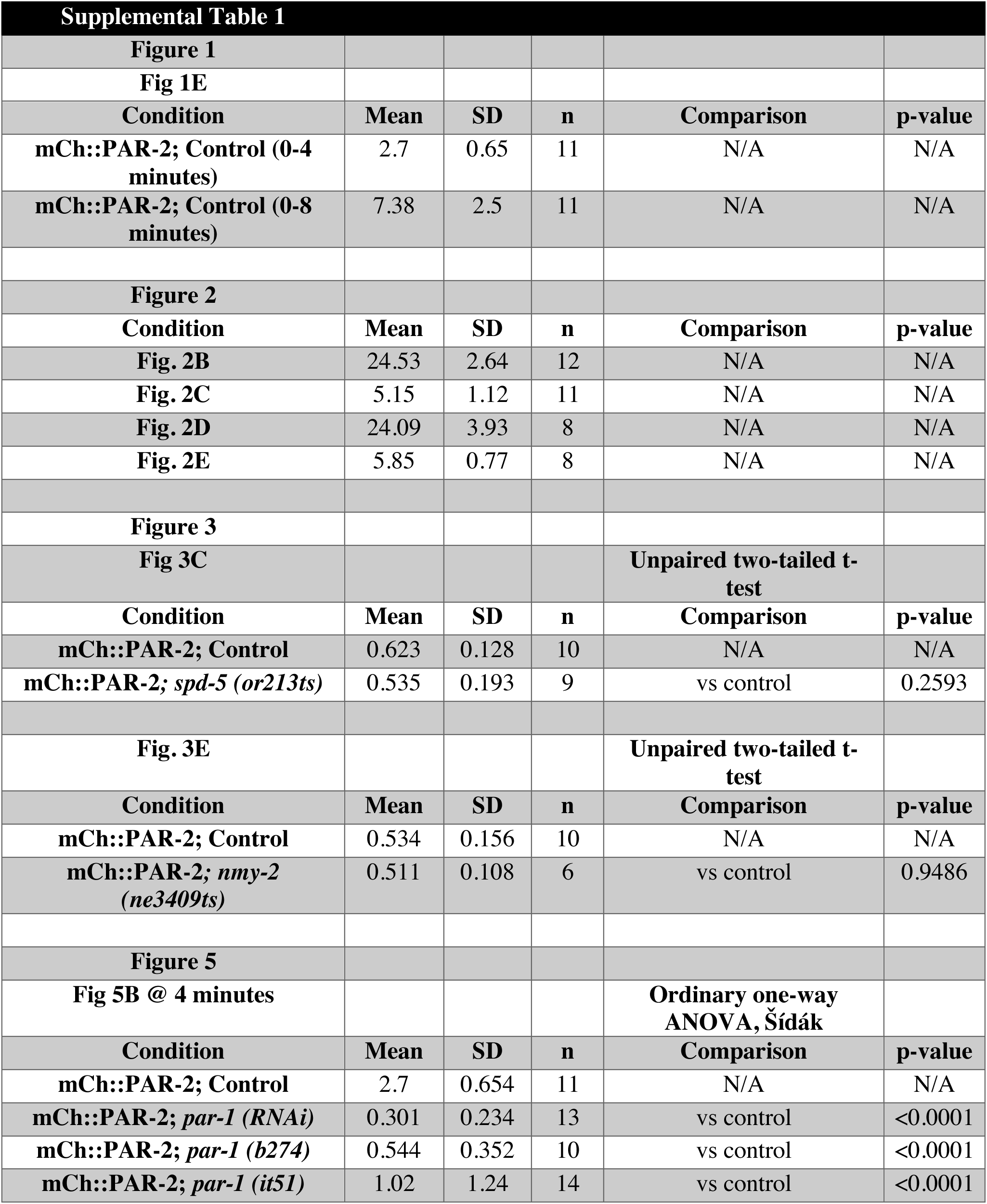
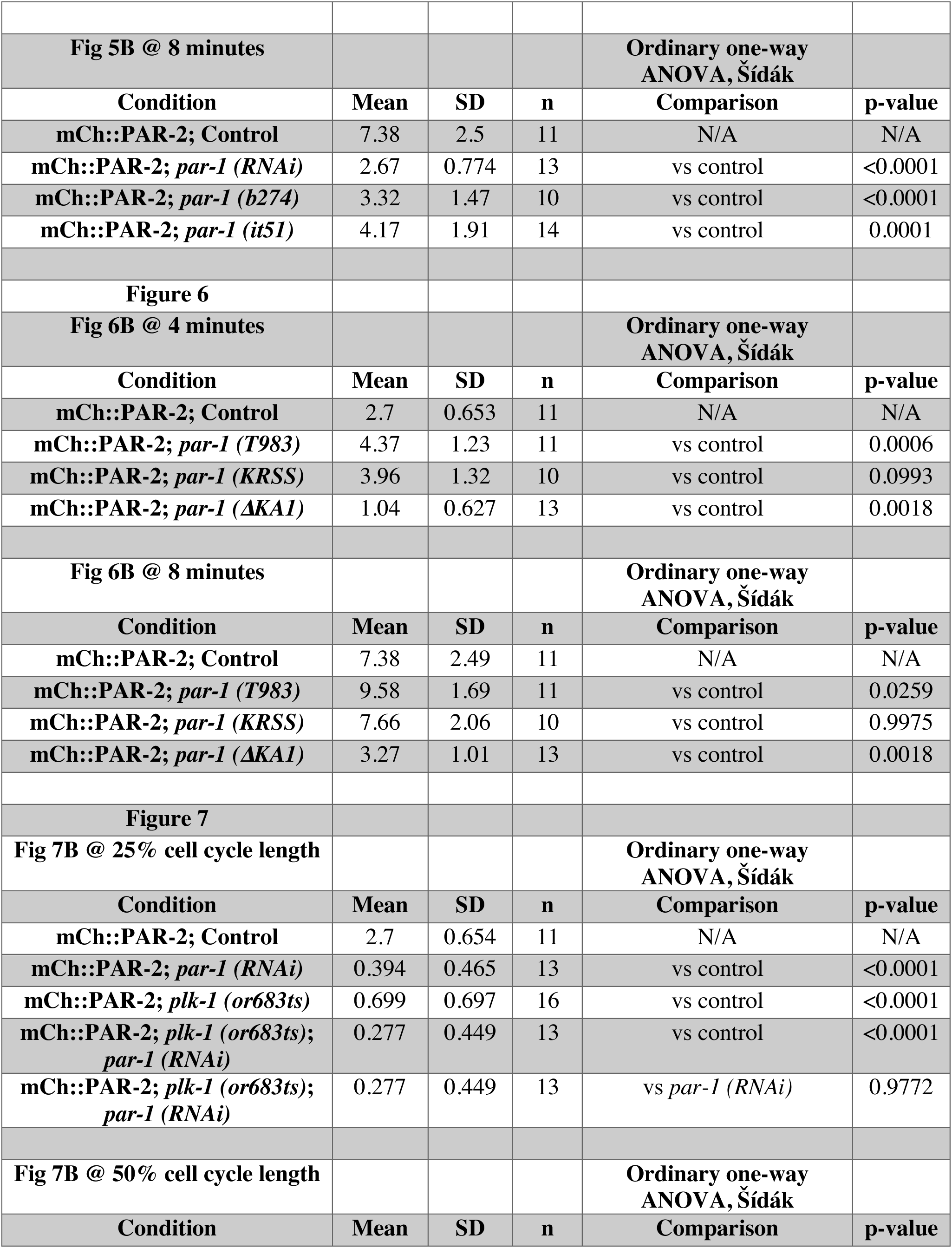
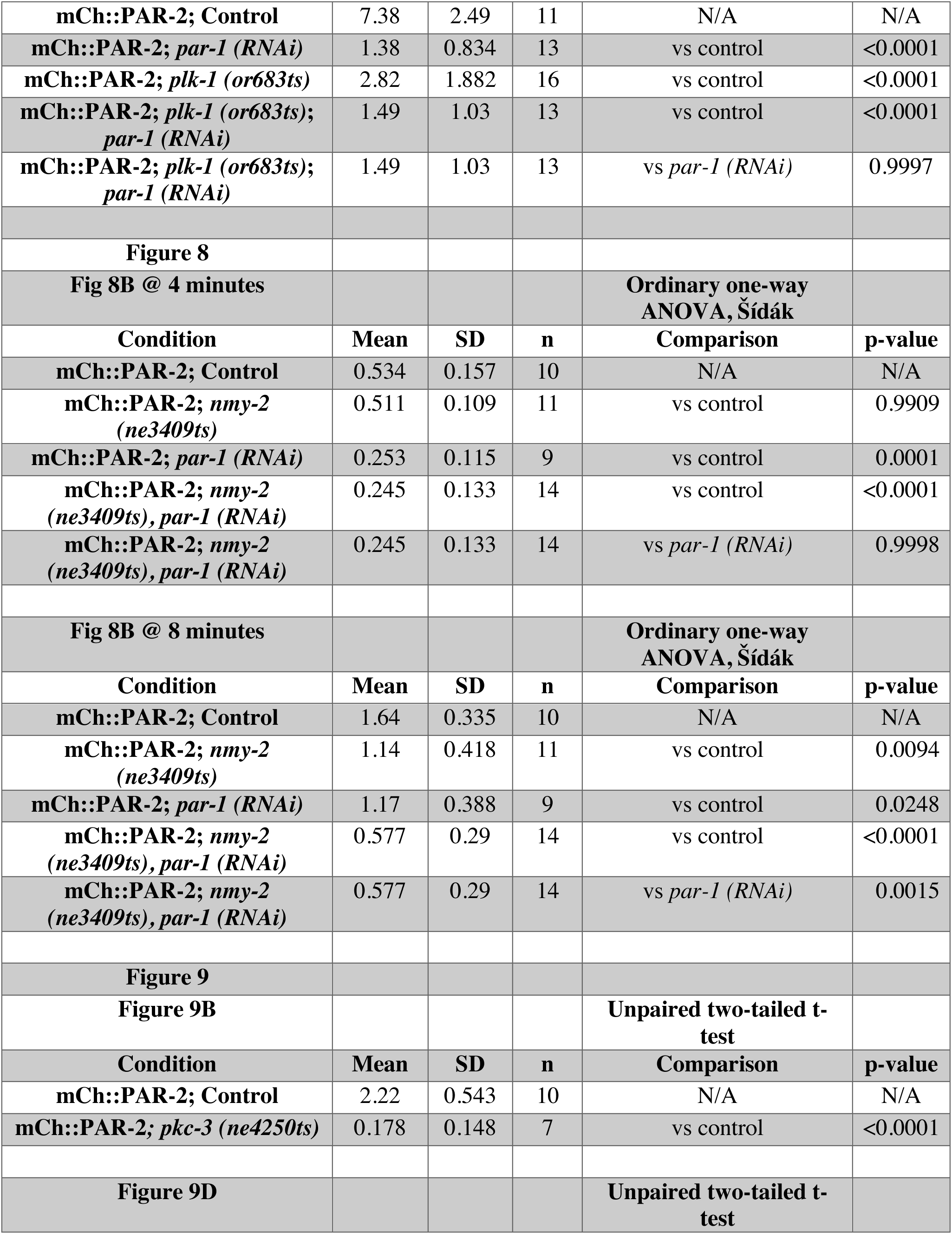
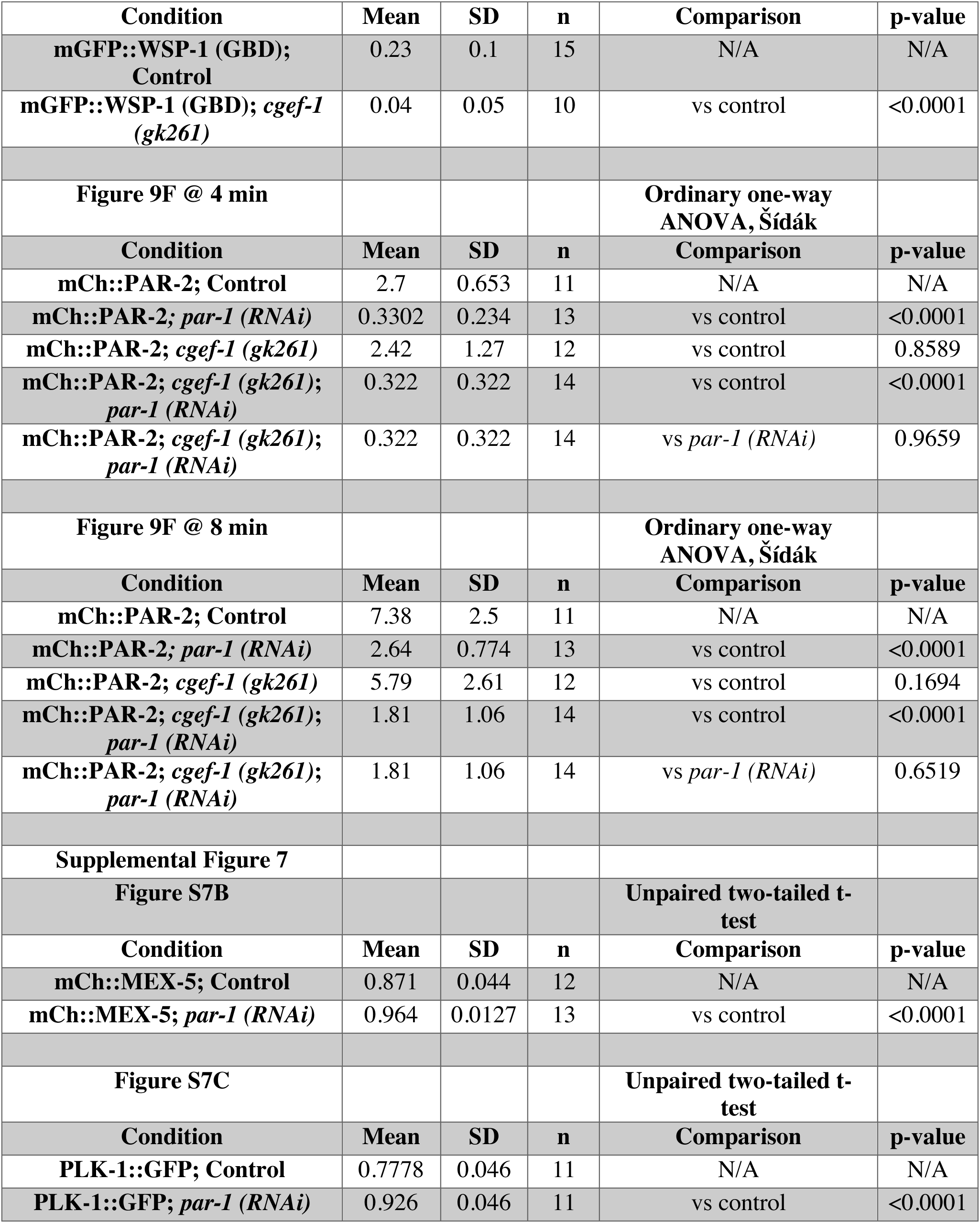
**–** Statistics corresponding to data presented in all figures and supplemental figures.

**Supplemental Movies -** All movies are oriented with anterior to the left. Original movies were taken at either 1 frame per 10 secs or 1 frame per 3 sec as indicated, but playback speed has been adjusted so that all play at same speed; timestamp is relative to end of cytokinesis (0:00).

**Supplemental_Video_1_ PAR2_PAR6** (10-second frame interval).

Movie corresponds to embryo used for Figure 1. Fluorescent images of an embryo expressing mCh::PAR-2 and GFP::PAR-6.

**Supplemental_Video_2_wild_type** (10-second frame interval).

Movie corresponds to embryo used for Figure 2A. DIC images of nuclear movement in the P1 cell in wild-type embryo.

**Supplemental_Video_3_ PAR2_AIR1** (10-second frame interval).

Movie corresponds to embryo used for Figure 2D. Confocal fluorescent images of embryos expressing mCh::PAR-2 and GFP::AIR-1 every minute from 0-4 minutes after the completion of cytokinesis.

**Supplemental_Video_4_ NMY2** (3-second frame interval).

Movie corresponds to embryo used for Figure 3A. Confocal fluorescent images of embryo expressing GFP::NMY-2 (Myosin) from a surface view.

**Supplemental_Video_5 _PAR2_wild_type** (10-second frame interval).

Movie corresponds to embryo used for control in Figure 5A. Confocal fluorescent images of mCh::PAR-2 in L4440 control embryo.

**Supplemental_Video_6 _PAR2_par-1(RNAi)** (10-second frame interval).

Movie corresponds to embryo used for *par-1(RNAi)* in Figure 5A. Confocal fluorescent images of mCh::PAR-2 in *par-1 (RNAi)* embryo.

## References

1. Arata, Y., Lee, J.-Y., Goldstein, B., & Sawa, H. (2010). Extracellular control of PAR protein localization during asymmetric cell division in the C. elegans embryo. Development, 137(19), 3337–3345. https://doi.org/10.1242/dev.054742

2. Bei, Y., Hogan, J., Berkowitz, L. A., Soto, M., Rocheleau, C. E., Pang, K. M., Collins, J., & Mello, C. C. (2002, Jul). SRC-1 and Wnt signaling act together to specify endoderm and to control cleavage orientation in early C. elegans embryos. Dev Cell, 3(1), 113–125. http://www.ncbi.nlm.nih.gov/pubmed/12110172

3. Benton, R., & St Johnston, D. (2003, Dec 12). Drosophila PAR-1 and 14-3-3 inhibit Bazooka/PAR-3 to establish complementary cortical domains in polarized cells. Cell, 115(6), 691–704. https://doi.org/10.1016/s0092-8674(03)00938-3

4. Boyd, L., Guo, S., Levitan, D., Stinchcomb, D. T., & Kemphues, K. J. (1996, Oct). PAR-2 is asymmetrically distributed and promotes association of P granules and PAR-1 with the cortex in C. elegans embryos. Development, 122(10), 3075–3084. https://doi.org/10.1242/dev.122.10.3075

5. Brenner, S. (1974, May). The genetics of Caenorhabditis elegans. Genetics, 77(1), 71–94. http://www.ncbi.nlm.nih.gov/pubmed/4366476

6. Cheeks, R. J., Canman, J. C., Gabriel, W. N., Meyer, N., Strome, S., & Goldstein, B. (2004, May 25). C. elegans PAR proteins function by mobilizing and stabilizing asymmetrically localized protein complexes. Curr Biol, 14(10), 851–862. https://doi.org/10.1016/j.cub.2004.05.022

7. Church, D. L., Guan, K. L., & Lambie, E. J. (1995, Aug). Three genes of the MAP kinase cascade, mek-2, mpk-1/sur-1 and let-60 ras, are required for meiotic cell cycle progression in Caenorhabditis elegans. Development, 121(8), 2525–2535. https://doi.org/10.1242/dev.121.8.2525

8. Cowan, C. R., & Hyman, A. A. (2004, Sep 2). Centrosomes direct cell polarity independently of microtubule assembly in C. elegans embryos. Nature, 431(7004), 92–96. https://doi.org/10.1038/nature02825

9. Cuenca, A. A., Schetter, A., Aceto, D., Kemphues, K., & Seydoux, G. (2003, Apr). Polarization of the C. elegans zygote proceeds via distinct establishment and maintenance phases. Development, 130(7), 1255–1265. https://doi.org/10.1242/dev.00284

10. Fievet, B. T., Rodriguez, J., Naganathan, S., Lee, C., Zeiser, E., Ishidate, T., Shirayama, M., Grill, S., & Ahringer, J. (2013, Jan). Systematic genetic interaction screens uncover cell polarity regulators and functional redundancy. Nat Cell Biol, 15(1), 103–112. https://doi.org/10.1038/ncb2639

11. Folkmann, A. W., & Seydoux, G. (2019, Mar 25). Spatial regulation of the polarity kinase PAR-1 by parallel inhibitory mechanisms. Development, 146(6). https://doi.org/10.1242/dev.171116

12. Goldstein, B. (1993, Aug). Establishment of gut fate in the E lineage of C. elegans: the roles of lineage-dependent mechanisms and cell interactions. Development, 118(4), 1267–1277. http://www.ncbi.nlm.nih.gov/pubmed/8269853

13. Goldstein, B. (1995, May). Cell contacts orient some cell division axes in the Caenorhabditis elegans embryo. J Cell Biol, 129(4), 1071–1080. http://www.ncbi.nlm.nih.gov/pubmed/7744956

14. Goldstein, B., & Macara, I. G. (2007, Nov). The PAR proteins: fundamental players in animal cell polarization. Dev Cell, 13(5), 609–622. https://doi.org/10.1016/j.devcel.2007.10.007

15. Griffin, E. E., Odde, D. J., & Seydoux, G. (2011, Sep 16). Regulation of the MEX-5 gradient by a spatially segregated kinase/phosphatase cycle. Cell, 146(6), 955–968. https://doi.org/10.1016/j.cell.2011.08.012

16. Guo, S., & Kemphues, K. J. (1995, May 19). par-1, a gene required for establishing polarity in C. elegans embryos, encodes a putative Ser/Thr kinase that is asymmetrically distributed. Cell, 81(4), 611–620. https://doi.org/10.1016/0092-8674(95)90082-9

17. Hamill, D. R., Severson, A. F., Carter, J. C., & Bowerman, B. (2002, Nov). Centrosome maturation and mitotic spindle assembly in C. elegans require SPD-5, a protein with multiple coiled-coil domains. Dev Cell, 3(5), 673–684. https://doi.org/10.1016/s1534-5807(02)00327-1

18. Hao, Y., Boyd, L., & Seydoux, G. (2006, Feb). Stabilization of cell polarity by the C. elegans RING protein PAR-2. Dev Cell, 10(2), 199–208. https://doi.org/10.1016/j.devcel.2005.12.015

19. Hurov, J. B., Watkins, J. L., & Piwnica-Worms, H. (2004, Apr 20). Atypical PKC phosphorylates PAR-1 kinases to regulate localization and activity. Curr Biol, 14(8), 736–741. https://doi.org/10.1016/j.cub.2004.04.007

20. Kamath, R. S., Fraser, A. G., Dong, Y., Poulin, G., Durbin, R., Gotta, M., Kanapin, A., Le Bot, N., Moreno, S., Sohrmann, M., Welchman, D. P., Zipperlen, P., & Ahringer, J. (2003, Jan 16). Systematic functional analysis of the Caenorhabditis elegans genome using RNAi. Nature, 421(6920), 231–237. https://doi.org/10.1038/nature01278

21. Kemphues, K. J., Priess, J. R., Morton, D. G., & Cheng, N. S. (1988, Feb 12). Identification of genes required for cytoplasmic localization in early C. elegans embryos. Cell, 52(3), 311–320. https://doi.org/10.1016/s0092-8674(88)80024-2

22. Kim, A. J., & Griffin, E. E. (2020). PLK-1 Regulation of Asymmetric Cell Division in the Early *C. elegans* Embryo. Front Cell Dev Biol, 8, 632253. https://doi.org/10.3389/fcell.2020.632253

23. Klinkert, K., Levernier, N., Gross, P., Gentili, C., von Tobel, L., Pierron, M., Busso, C., Herrman, S., Grill, S. W., Kruse, K., & Gönczy, P. (2019, Feb 26). Aurora A depletion reveals centrosome-independent polarization mechanism in Caenorhabditis elegans. Elife, 8. https://doi.org/10.7554/eLife.44552

24. Kumfer, K. T., Cook, S. J., Squirrell, J. M., Eliceiri, K. W., Peel, N., O’Connell, K. F., & White, J. G. (2010, Jan 15). CGEF-1 and CHIN-1 regulate CDC-42 activity during asymmetric division in the Caenorhabditis elegans embryo. Mol Biol Cell, 21(2), 266–277. https://doi.org/10.1091/mbc.e09-01-0060

25. Liu, J., Maduzia, L. L., Shirayama, M., & Mello, C. C. (2010, Mar 15). NMY-2 maintains cellular asymmetry and cell boundaries, and promotes a SRC-dependent asymmetric cell division. Dev Biol, 339(2), 366–373. https://doi.org/10.1016/j.ydbio.2009.12.041

26. Motegi, F., Zonies, S., Hao, Y., Cuenca, A. A., Griffin, E., & Seydoux, G. (2011, Oct 09). Microtubules induce self-organization of polarized PAR domains in Caenorhabditis elegans zygotes. Nat Cell Biol, 13(11), 1361–1367. https://doi.org/10.1038/ncb2354

27. Munro, E., Nance, J., & Priess, J. R. (2004, Sep). Cortical flows powered by asymmetrical contraction transport PAR proteins to establish and maintain anterior-posterior polarity in the early C. elegans embryo. Dev Cell, 7(3), 413–424. https://doi.org/10.1016/j.devcel.2004.08.001

28. Nishi, Y., Rogers, E., Robertson, S. M., & Lin, R. (2008, Feb). Polo kinases regulate C. elegans embryonic polarity via binding to DYRK2-primed MEX-5 and MEX-6. Development, 135(4), 687–697. https://doi.org/10.1242/dev.013425

29. O’Rourke, S. M., Carter, C., Carter, L., Christensen, S. N., Jones, M. P., Nash, B., Price, M. H., Turnbull, D. W., Garner, A. R., Hamill, D. R., Osterberg, V. R., Lyczak, R., Madison, E. E., Nguyen, M. H., Sandberg, N. A., Sedghi, N., Willis, J. H., Yochem, J., Johnson, E. A., & Bowerman, B. (2011, Mar 1). A survey of new temperature-sensitive, embryonic-lethal mutations in C. elegans: 24 alleles of thirteen genes. PLoS One, 6(3), e16644. https://doi.org/10.1371/journal.pone.0016644

30. Pickett, M. A., Naturale, V. F., & Feldman, J. L. (2019, Oct 6). A Polarizing Issue: Diversity in the Mechanisms Underlying Apico-Basolateral Polarization In Vivo. Annu Rev Cell Dev Biol, 35, 285–308. https://doi.org/10.1146/annurev-cellbio-100818-125134

31. Portier, N., Audhya, A., Maddox, P. S., Green, R. A., Dammermann, A., Desai, A., & Oegema, K. (2007, Apr). A microtubule-independent role for centrosomes and aurora a in nuclear envelope breakdown. Dev Cell, 12(4), 515–529. https://doi.org/10.1016/j.devcel.2007.01.019

32. Reich, J. D., Hubatsch, L., Illukkumbura, R., Peglion, F., Bland, T., Hirani, N., & Goehring, N. W. (2019, Jun 17). Regulated Activation of the PAR Polarity Network Ensures a Timely and Specific Response to Spatial Cues. Curr Biol, 29(12), 1911–1923.e1915. https://doi.org/10.1016/j.cub.2019.04.058

33. Rodriguez, J., Peglion, F., Martin, J., Hubatsch, L., Reich, J., Hirani, N., Gubieda, A. G., Roffey, J., Fernandes, A. R., St Johnston, D., Ahringer, J., & Goehring, N. W. (2017, Aug 21). aPKC Cycles between Functionally Distinct PAR Protein Assemblies to Drive Cell Polarity. Dev Cell, 42(4), 400–415.e409. https://doi.org/10.1016/j.devcel.2017.07.007

34. Rose, L., & Gonczy, P. (2014). Polarity establishment, asymmetric division and segregation of fate determinants in early C. elegans embryos. WormBook, 1-43. https://doi.org/10.1895/wormbook.1.30.2

35. Schonegg, S., Hyman, A. A., & Wood, W. B. (2014, Jun). Timing and mechanism of the initial cue establishing handed left–right asymmetry in Caenorhabditis elegans embryos. Genesis, 52(6), 572–580. https://doi.org/10.1002/dvg.22749

36. Schubert, C. M., Lin, R., de Vries, C. J., Plasterk, R. H., & Priess, J. R. (2000, Apr). MEX-5 and MEX-6 function to establish soma/germline asymmetry in early C. elegans embryos. Mol Cell, 5(4), 671–682. https://doi.org/10.1016/s1097-2765(00)80246-4

37. Seirin-Lee, S., Gaffney, E. A., & Dawes, A. T. (2020, Sep 5). CDC-42 Interactions with Par Proteins Are Critical for Proper Patterning in Polarization. Cells, 9(9). https://doi.org/10.3390/cells9092036

38. Sunchu, B., & Cabernard, C. (2020, Jun 29). Principles and mechanisms of asymmetric cell division. Development, 147(13). https://doi.org/10.1242/dev.167650

39. Tabuse, Y., Izumi, Y., Piano, F., Kemphues, K. J., Miwa, J., & Ohno, S. (1998, Sep). Atypical protein kinase C cooperates with PAR-3 to establish embryonic polarity in Caenorhabditis elegans. Development, 125(18), 3607–3614. http://www.ncbi.nlm.nih.gov/pubmed/9716526

40. Timmons, L., & Fire, A. (1998, Oct 29). Specific interference by ingested dsRNA. Nature, 395(6705), 854. https://doi.org/10.1038/27579

41. Venkei, Z. G., & Yamashita, Y. M. (2018, Nov 5). Emerging mechanisms of asymmetric stem cell division. J Cell Biol, 217(11), 3785–3795. https://doi.org/10.1083/jcb.201807037

42. Watts, J. L., Etemad-Moghadam, B., Guo, S., Boyd, L., Draper, B. W., Mello, C. C., Priess, J. R., & Kemphues, K. J. (1996, Oct). par-6, a gene involved in the establishment of asymmetry in early C. elegans embryos, mediates the asymmetric localization of PAR-3. Development, 122(10), 3133–3140. https://doi.org/10.1242/dev.122.10.3133

43. Zhao, P., Teng, X., Tantirimudalige, S. N., Nishikawa, M., Wohland, T., Toyama, Y., & Motegi, F. (2019, Mar 11). Aurora-A Breaks Symmetry in Contractile Actomyosin Networks Independently of Its Role in Centrosome Maturation. Dev Cell, 48(5), 631–645.e636. https://doi.org/10.1016/j.devcel.2019.02.012

44. Zonies, S., Motegi, F., Hao, Y., & Seydoux, G. (2010, May). Symmetry breaking and polarization of the C. elegans zygote by the polarity protein PAR-2. Development, 137(10), 1669–1677. https://doi.org/10.1242/dev.045823

